# Evolutionary Insights from the Mitochondrial Genome of *Oikopleura dioica*: Sequencing Challenges, RNA Editing, Gene Transfers to the Nucleus, and tRNA Loss

**DOI:** 10.1101/2024.05.09.593300

**Authors:** Yael Klirs, Maria Novosolov, Carmela Gissi, Rade Garic, Tal Pupko, Thomas Stach, Dorothée Huchon

## Abstract

Sequencing the mitochondrial genome of the tunicate *Oikopleura dioica* is a challenging task due to the presence of long poly-A/T homopolymer stretches, which impair sequencing and assembly. Here, we report on the sequencing and annotation of the majority of the mitochondrial genome of *O. dioica* by means of combining several DNA and amplicon reads obtained by Illumina and MinIon Oxford Nanopore Technologies (ONT) with public RNA sequences. We document extensive RNA editing, since all homopolymer stretches present in the mitochondrial DNA correspond to 6U-regions in the mitochondrial RNA. Out of the 13 canonical protein-coding genes, we were able to detect eight, plus an unassigned ORF that lacked sequence similarity to canonical mitochondrial protein-coding genes. We show that the *nad3* gene has been transferred to the nucleus and acquired a mitochondria-targeting signal. In addition to two very short rRNAs, we could only identify a single tRNA (tRNA-Met), suggesting multiple losses of tRNA genes, supported by a corresponding loss of mitochondrial aminoacyl-tRNA synthetases in the nuclear genome. Based on the eight canonical protein-coding genes identified, we reconstructed maximum likelihood and Bayesian phylogenetic trees and inferred an extreme evolutionary rate of this mitochondrial genome. The phylogenetic position of appendicularians among tunicates, however, could not be accurately determined.

**Significance:** Sequencing and annotating the mitochondrial genome of fast-evolving organisms is difficult because they often present unusual characteristics. The tunicate *Oikopleura dioica* is a model species for understanding tunicate and chordate genome evolution. However, no complete annotated mitochondrial genome for this species has been published to date. Here, we determined the major part of the mitochondrial genome of *O. dioica*. Our results indicate the presence of highly modified rRNA genes and the absence of all tRNAs except tRNA-Met. Moreover, we show that the mitochondrial genome undergoes editing at the RNA level. Our study demonstrates that utilizing a combination of public RNA data and DNA from long-and short-read sequencing platforms significantly improves our ability to study mitochondrial genomes with atypical characteristics.

## Introduction

Most mitochondrial genomes of the Bilateria are circular DNA molecules of approximately 14-19 kb in length, characterized by the absence of introns and the presence of only short intergenic regions (Lavrov and Pett 2016). These genomes typically contain 37 genes: 13 protein-coding genes: ATP synthase subunits 6 and 8 (*atp6* and *atp8*), cytochrome oxidase subunits (*cox1, cox2, cox3*), cytochrome b (*cytb*), and dehydrogenase subunits (*nad1, nad2, nad3, nad4, nad5, nad6*, and *nad4L*); two ribosomal genes (rRNA): small-subunit (*rns*) and large-subunit (*rnl*) and ∼22 tRNAs (Boore 1999; Gissi et al. 2008; Rubinstein et al. 2013). However, bilaterian mitochondrial genomes are substantially diverse in their structure, organization, evolutionary rate, and genetic code (Lavrov 2007).

Tunicate mitochondrial genomes are fast evolving, both in terms of substitutions (Singh et al. 2009) and genome rearrangements (Gissi et al. 2008; Stach et al. 2010). As a result of this high rearrangement rate, there is no standard tunicate mitochondrial gene order. In addition, the ascidian mitochondrial genetic code differs from the vertebrate one. Of note, the “ascidian” genetic code, which was initially determined based on the mitochondrial sequences of *Halocynthia roretzi* and *Pyura stolonifera* (Durrheim et al. 1993; Yokobori et al. 1993), seems to be valid for all other tunicate classes (Pichon et al. 2020).

The subphylum Tunicata is divided into three classes: Ascidiacea, Thaliacea, and Appendicularia. Appendicularia is a key class for understanding tunicate and chordate genome evolution, as it seems to be the sister-clade to all other tunicates. *Oikopleura dioica* is a frequent model tunicate for the class Appendicularia in developmental biology, evolutionary biology, and comparative genomics research (Delsuc et al. 2006; Nishida 2008). Despite the importance of this clade, no complete annotated appendicularian mitochondrial genome is currently available in public databases. One non-annotated mitochondrial genome of *O. dioica* from Okinawa Japan is available (accession OU015571; 9,225 bp; (Bliznina, et al. 2021)).

The mitochondrial genome of *O. dioica* from Norway has been partially reconstructed by Denoeud et al. (2010) from genomic DNA and EST data. However, the obtained mitochondrial contigs of this genome are not publicly available. This genome has been inferred to harbor long poly-A/T stretches interrupting the open reading frames (ORFs) and it has been suggested that these interrupted ORFs are corrected by RNA editing (Denoeud et al. 2010). Such homopolymer stretches cannot be sequenced using Sanger sequencing or Illumina short-read sequencing, since they are error hot spots for the DNA polymerases used in these sequencing techniques. Thus, together with the difficulty in isolation and identification of appendicularian species, this peculiarity explains the paucity of publicly available mitochondrial data on *O. dioica*. Annotation of the resulting genome is also expected to be challenging, due to RNA editing. RNA editing refers to modifications such as insertions, deletions, and base substitutions of RNA transcripts. As a result, the RNA and computationally inferred protein sequences differ from the DNA template (Gott and Emeson 2000). To complicate matters, in *O. dioica* RNA editing seems to be prevalent in those regions that are hard to sequence, i.e., the poly-A/T stretches. These stretches appear as 6U-sites in mitochondrial RNA sequences, regardless of the number of T/A at the DNA level (Denoeud et al. 2010).

The phylogenetic analyses performed by Denoeud et al. (2010) concluded that *O. dioica* mitochondrial protein-coding genes are very fast evolving, even when compared to the already high rate of other tunicate mitochondrial genomes. Using a combination of PCR and EST mapping approaches, they determined the order of nine out of the 13 mitochondrial protein-coding genes: *cox3, nad1, cox1, cytb,* a putative *nad2*, *atp6, cox2, nad5*, along a large mitochondrial contig, and *nad4* on a separate contig. They did not identify any tRNA or rRNA genes. Unfortunately, the mitochondrial contigs obtained by Denoeud et al. (2010) have not been deposited in public databases and only the raw ESTs data are available in the Oikobase database (Danks et al. 2012).

A draft mitochondrial genome scaffold of *O. dioica* from Okinawa (Japan) has been published by Bliznina et al. (2021) (accession OU015571; 9,225 bp), with the annotation reported only in the publication (Figure 7 of their article). The annotation obtained is, however, preliminary as several genes contain stop codons and frameshifts. Unlike in Denoeud et al. (2010), the mitochondrial genome is presented as a circular molecule that encompasses seven mitochondrial genes (Bliznina et al. 2021). The mitochondrial gene order resembles that from Figure S4 from Denoeud et al. (2010). However, some differences exist. First, the *nad2* and *nad5* genes are missing from the annotation of Bliznina et al. (2021). Second, the *rnl* and four tRNAs: tRNA-Met, tRNA-Trp, tRNA-Ala, and tRNA-His are annotated. Third, they identified a LAGLIDADG homing endonuclease in the middle of the *cox1* gene (Figure 7 from Bliznina et al. 2021). It should be noted that *O. dioica* from Okinawa most likely represents a different species from *O. dioica* from Norway (Masunaga et al. 2022; Plessy et al. 2024).

Here, we determined the major part of the mitochondrial genome sequence of *O. dioica* from Norway, its gene content, and its order. We observed a substantial gene loss compared to other metazoan mitochondrial genomes and provide evidence for extensive RNA editing.

## Results

### Sequencing the Mitochondrial Genome

DNA extracted from *O. dioica* individuals from Norway was sent for Illumina sequencing. Since it was impossible to assemble these reads due to the prevalence of homopolymers (poly-A or poly-T stretches), we next performed long-read sequencing using Oxford Nanopore Technology (ONT) sequencing. However, in this case, the flow-cell pores became blocked within an hour (Minion sequencing was performed twice by different companies), resulting in too little data for a reliable assembly. Therefore, we PCR-amplified the mitochondrial genome in 12 fragments (supplementary Fig. S1, Supplementary Material online) and sequenced all barcoded fragments using a single ONT flow cell. Primers were designed based on the *O. dioica* EST and RNA data available in public databases (see Materials and Methods and supplementary table S1, Supplementary Material online). We successfully assembled each PCR fragment (except one) and, using the Illumina data, we obtained a single contig. Unfortunately, we could not circularize the genome because we were unable to assemble one fragment completely. The length of the *O. dioica* mitochondrial contig we assembled is 18,390 bp (Figure 1). This genome size is larger than the average of other tunicates, 14,808 ± 821 bp, and compatible with the PCR prediction of a ∼19,000 bp genome (supplementary table S1, Supplementary Material online).

**Figure 1.**
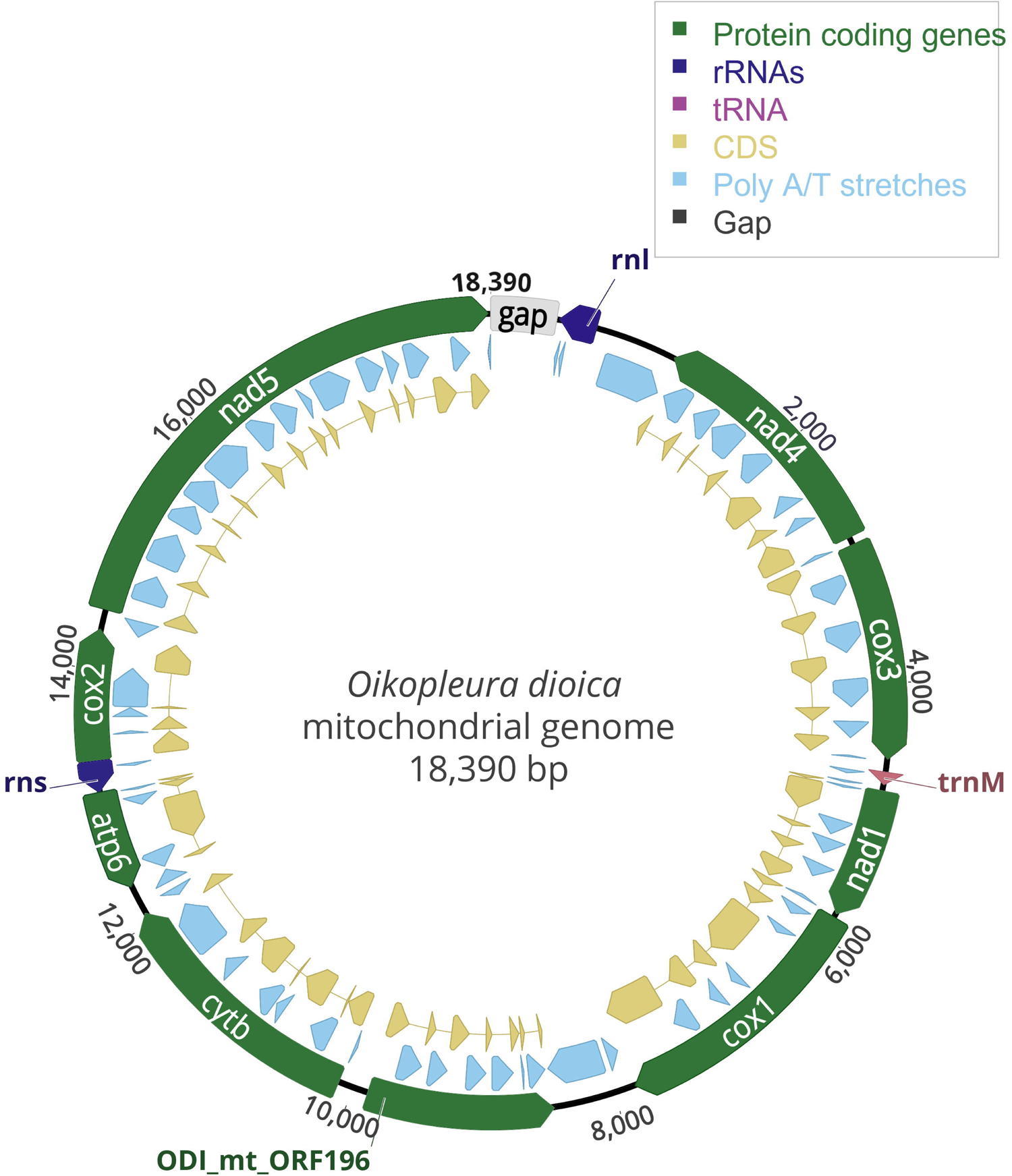
*Oikopleura dioica* mitochondrial genome map. Protein-coding genes are in green, rRNA genes (*rns*, *rnl*) in dark blue, tRNA-Met (UAU) in pink, coding sequences (CDS) in yellow, and poly-A/T stretches in light blue. The genes are pointing in the direction of their transcriptional orientation. A non-sequenced region, indicated as “gap”, is in gray.

Multiple instances of poly-A/T stretches were found in the assembled contig (Figure 1, supplementary Fig. S2, Supplementary Material online). The exact number of either A/T repeats within poly-A/T regions could not be accurately determined. These stretches are too long to be fully contained in the short Illumina sequences (Ari and Arikan 2016), while ONT basecallers do not accurately determine the length of homopolymers (Wick et al. 2019). Moreover, the length of homopolymers has been correctly resolved up to a length of 11Lbp in an R10.4 flow cell (Sereika et al. 2022), while in an R9.4.1 flow cell, a good accuracy (70% errorless) was observed only in homopolymers up to 5 bp (Delahaye and Nicolas 2021). However, in our case, the length of the 65 poly-A/T stretches found in the mitochondrial sequence ranged from 2 to 509 bp, with an average of 143 ± 103.5 bp (Materials and Methods, supplementary table S2, Supplementary Material online), which is considerably larger. Moreover, it has been observed that regions harboring inverted repeats are poorly sequenced using ONT (Spealman et al. 2020). When in a PCR-fragment a poly-A region is followed by a poly-T region, these two homopolymers can form structures similar to inverted repeats, thus disrupting ONT sequencing. In addition, Taq polymerase slippage errors can further contribute to uncertainty associated with the length of long homopolymers. All these factors contribute to heterogeneity among ONT reads that map to the same mitochondrial locus harboring a poly-A/T region. One such example is shown in Figure 2. Note that these stretches are sometimes imperfect, i.e., there may be a few G residues within a poly-A stretch or a few C residues with a poly-T stretch (Figure 2). These G and C residues are unlikely to reflect ONT sequencing errors, as they could sometimes be verified using Illumina reads. In total, these poly-A/T stretches span ∼9,200 bp. This suggests that while the total mitochondrial genome size is longer than in other tunicates, the mitochondrial genome without the poly-A/T stretches is, in fact, shorter.

**Figure 2.**
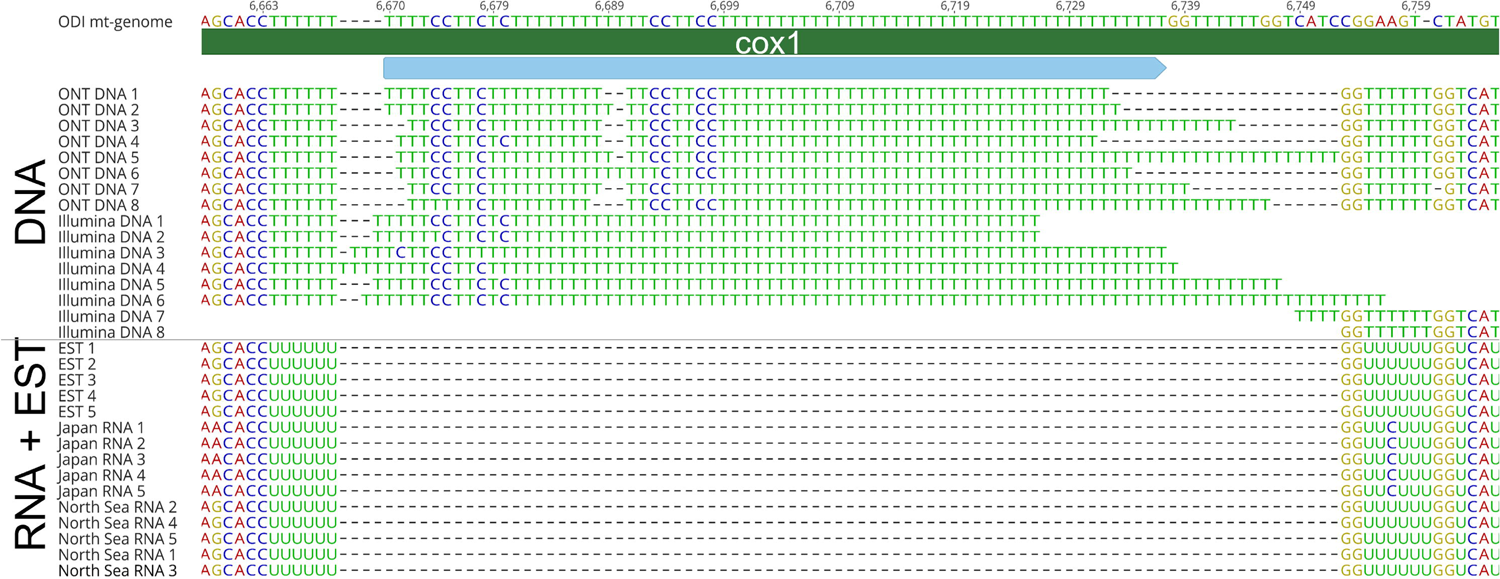
A typical poly-A/T region in the *O. doica cox1* gene. The reference sequence represents the assembled mitochondrial genome. The sequence of part of the *cox1* gene is indicated in dark green. Above it are the nucleotide and amino-acid sequences of the reference sequence. The poly-A/T region is in light blue. Reads from five different sequencing sources mapped to this region are shown below the reference sequence: our ONT data, our Illumina reads, public EST data from Norway, public RNA reads from Japan, and public RNA reads from the North Sea (see Supplementary table S3A, Supplementary Material online for accession numbers). Each line represents a different read. Note that the different ONT reads do not have the same length, exemplifying the fact that the length of the poly-T region could not be accurately determined. The EST and RNA reads lack a long poly-U region. Rather, the poly-T region corresponds to 6U-bases. This difference (long poly-T stretch in DNA *versus* 6U in the EST and RNA reads) provides evidence of editing at the RNA level. Illumina reads can map to each side of the poly-T stretch, but because poly-T stretches can be longer than the read length, we were unable to assemble the entire region using Illumina only or to determine the length of the poly-T region.

### Evidence of rRNA Editing

A total of 65 poly-A/T sites were found in the assembled *O. dioica* mitochondrial genome. By mapping (1) the mitochondrial transcripts identified from the EST from Norway, and (2) the RNA contigs assembled from publicly available RNA reads from mainland Japan (Wang et al. 2015) and the North Sea (Raghavan et al. 2023) onto the DNA sequence assembled from the ONT reads, we were able to confirm the editing of 59 of these 65 poly-A/T sites to 6U-sites (see an example in Figure 2 and supplementary table S2, Supplementary Material online). Of note, we performed a barcoding analysis that supports the hypothesis that our DNA sample, the EST data, and the North Sea RNA reads originated from the same species, while the RNA reads from mainland Japan likely originated from a different species (see Materials and Methods, Masunaga et al. 2022; Plessy et al. 2024). For the remaining six sites, no direct RNA evidence for editing was obtained as we could not detect RNA reads or ESTs that mapped to the corresponding region. These poly-A/T stretches are present both in coding and noncoding mitochondrial DNA regions. Specifically, we observed that exactly six consecutive U residues (6U-sites) in the RNA corresponded to a poly-T stretch in the DNA sequence. These results suggest that the longer U stretches in the pre-mRNA are edited, resulting in shorter stretches of 6U. The exact nature of this RNA editing is unknown. It is, however, remarkable that protein-coding genes as well as the two rRNAs and the tRNA-Met only include poly-T regions and not poly-A regions. When poly-A regions are found on the direct strand, this indicates that the genes are encoded on the reverse strand. We detected a few instances of EST reads that include a poly-U stretch, i.e., before the excision of poly-U to 6U. These observations suggest that the shortening of the poly-U stretch, which is necessary to recover the reading frame, occurs at the RNA level co-transcriptionally or very soon after the synthesis of the precursor transcript.

Interestingly we noticed that while poly-T regions are present within all genes, the genes often end with a poly-A. The poly-A ending is part of the TAA stop codon in most protein-coding genes, except for *cox1*, which has a TAG stop codon (Table 1). The contigs assembled from RNA reads support the view that these poly-A stretches are reduced to 6A in the mRNA. Similarly, we noticed that a poly-A stretch is often present in the DNA before the start of a protein-coding gene (Figure 1). No conserved sequence motif could be identified around the poly-A/T stretches. The only major departure we could detect around the poly-A/T stretches was the absence of cytosine just after the 6T site. No other signal could be identified (Supplementary Fig. S3, Supplementary Material online).

**Table 1.** *Oikopleura dioica* genes, position, length, direction (F-forward, R-reverse), start and stop codons, and comparison to the minimal, maximal, and average length of orthologous genes in 38 tunicate sequences (supplementary table S4, Supplementary Material online).

| Gene | Type | Position |  | Length [bp] |  | Direction | Codon |  | Length range in tunicates [bp] |  | Average length in tunicates [bp] |
| --- | --- | --- | --- | --- | --- | --- | --- | --- | --- | --- | --- |
|  |  | start | end | gene | CDS |  | start | stop | min | max |  |
| *rnl* | rRNA | 13 | 303 | 291 | 242 | R |  |  | 921 | 1,299 | 1,105 |
| *nad4* | gene | 934 | 2,847 | 1,914 | 876 | R | ATC | TAA | 1,209 | 1,401 | 1,321 |
| *cox3* | gene | 2,906 | 4,574 | 1,669 | 870 | F | ATT | TAA | 768 | 795 | 786 |
| *tRNA-Met* | tRNA | 4,671 | 4,763 | 93 | 69 | F |  |  | 61 | 76 | 66 |
| *nad1* | gene | 4,805 | 5,800 | 996 | 615 | F | TTG | TAA | 853 | 1,035 | 914 |
| *cox1* | gene | 5,836 | 7,834 | 1,999 | 1,512 | F | ATT | TAG | 1,482 | 1,575 | 1,546 |
| *ODI_mt_ORF196* | gene | 8,462 | 9,887 | 1,426 | 591 | R | TTG | TAA |  |  |  |
| *cytb* | gene | 10,115 | 12,085 | 1,971 | 993 | F | GTG | TAA | 1,035 | 1,122 | 1,093 |
| *atp6* | gene | 12,312 | 13,063 | 752 | 516 | R | ATT | TAA | 492 | 642 | 619 |
| *rns* | rRNA | 13,064 | 13,295 | 232 | 215 | R |  |  | 596 | 984 | 696 |
| *cox2* | gene | 13,305 | 14,301 | 997 | 582 | F | ATT | TAA | 651 | 942 | 689 |
| *nad5* | gene | 14,456 | 18,371 | 3,916 | 1,248 | F | ATA | TAA | 1,458 | 1,788 | 1,695 |

### *Oikopleura dioica* Mitochondrial Genome Sequence and Structure

Figure 1 presents the assembled mitochondrial genome of *O. dioica*. We could not determine the sequence in the region labeled as “Gap” between the *nad5* and *rnl* genes, probably due to the presence of secondary structures that impaired sequencing. Running the PCR product of fragment 12 on a gel indicated that the length of the “Gap” region is approximately 400 bp (supplementary Fig. S1, supplementary table S1, Supplementary Material online). However, the fact that we could amplify this missing region using primers from both sides of the gap suggests that the genome is circular. Notably, although we could not assemble the entire fragment 12, the ONT-contigs we obtained allowed us to determine the end of the *nad5* gene and the *nad4*-*rnl* region, supporting the view that we had amplified the correct region of the mitochondrial genome. The circularity of the mitochondrial genome is in line with Bliznina et al. (2021).

### Gene Content and Gene Loss in *O. dioica* Mitochondrial Genome

Most tunicate mitochondrial genomes encompass 13 protein-coding genes, two rRNA genes, and 24 tRNA genes that are encoded on the same strand (Shenkar et al. 2016; Singh et al. 2009). Surprisingly, in the mitochondrial genome of *O. dioica* we could only identify eight canonical protein-coding genes: ATP synthase subunit 6, subunits 1-3 of cytochrome c oxidase (*cox*), cytochrome *b* (*cytb*), and NAD dehydrogenase (*nad*) 1, 4, and 5. In addition to the two rRNA genes encoding the small and large subunits, we were able to identify a single tRNA gene, tRNA-Met (Figure 1, Table 1). The genes *cox1, cox2, cox3, cytb, nad1, nad5*, and tRNA-Met (UAU) are encoded on the forward strand, while *atp6, nad4*, *rns*, *rnl,* and an unidentified protein-coding gene named *ODI_mt_ORF196* (see below), are encoded on the reverse strand (Table 1). No incomplete stop codons were found (Table 1). Moreover, we did not find any LAGLIDADG homing endonuclease ORF encoded within the *cox1* gene. These findings suggest the loss of five protein-coding genes: *atp8, nad 2, 3, 4L*, and *6* and, all tRNA genes except tRNA-Met.

An additional mitochondrial protein-coding gene was identified following a Blastn search of the obtained mitochondrial sequence against the EST data available in Oikobase (Danks et al. 2012). Notably, programs such as “Find ORFs” (implemented in Geneious Prime v.2023.2.1) are inapplicable for this mitochondrial genome, due to the extremely fast rate of evolution and the numerous poly-A/T stretches that disrupt the reading frames of protein-coding genes. This additional ORF is 1,426 bp long and does not show any significant sequence similarity to any known protein-coding gene present in the NCBI public database (i.e., based on Blastp searches). We thus termed it gene “*ODI_mt_ORF196*” based on its amino-acid length (Figure 1, Table 1). This protein was inferred to have six transmembrane domains and, based on its position, it corresponds to the putative *nad2* gene identified by Denoeud et al. (2010). An Alpha fold-based prediction of this gene (supplementary Fig. S4, Supplementary Material online) was used as query against multiple protein structure databases: AlphaFold/UniProt50 v4, AlphaFold/Swiss-Prot v4, AlphaFold/Proteome v4, CATH50 4.3.0, MGnify-ESM30 v1, PDB100 2201222, GMGCL 2204 from Foldseek (van Kempen et al. 2024). No significant structural hits were obtained.

We identified the two rRNA genes, *rns* and *rnl,* by searching for highly transcribed regions along unannotated regions of the assembled genome. Two non-protein-coding regions were found to be highly expressed based on Illumina RNA read mapping (supplementary Fig. S5, Supplementary Material online). Further support for the transcription of these regions lies in their presence in the EST data. Specifically, among these two highly expressed regions, the longer one, which resides between the “gap region” and the *nad4* gene, was assumed to be the *rnl* gene. This annotation is supported by RNA structure similarities between the putative *rnl* and the structure model profile of this gene, as implemented in Mitos2 (Donath et al. 2019). The second one, located between the *atp6* and the *cox2* genes, was annotated as *rns*, although no structural similarity was found between this sequence and the *rns* structure profile in Mitos2. Both the *rnl* and *rns* genes were found to be extremely short: 291 and 232 bp, respectively, compared to 1,129 and 660 bp in the model tunicate *Ciona intestinalis* (see also Table 1).

We were able to identify a single tRNA gene, tRNA-Met (UAU). This tRNA presents a typical secondary cloverleaf structure (supplementary Fig. S6, Supplementary Material online), and UAU anti-codon, corresponding to a methionine codon according to the ascidian mitochondrial genetic code. Unlike in Bliznina et al. (2021), no other tRNAs were identified.

### Aminoacyl-tRNA Synthetase (aaRS) Profiling

We used hidden Markov models (HMMs) to identify nuclear-encoded aaRSs in published sequences of *O. dioica* and a few other chordate representatives. We found the presence of all cytosolic aaRSs in published proteomes of *O. dioica* (accessions: GCA_000209535, Norway and GCA_907165135, Okinawa) (Figure 3). However, in contrast to the other chordates investigated, *O. dioica* is inferred to have lost most of its mitochondrial aaRSs except for aaRS-Gly (G), aaRS-Phe (F), and aaRS-Met (M). Similarly, the mitochondrial aaRS-Val (V) is absent in the tunicates *C. intestinalis* (GCA_000224145) and *Styela clava* (GCF_000001405), and the mitochondrial aaRS-Lys (K) and aaRS-Thr (T) are absent in all the species included in this comparison. Furthermore, *O. dioica* encodes two copies of aaRS-Gly (one cytosolic and the other mitochondrial), while the other chordates encode only a single copy that functions both in the cytosol and the mitochondrion, suggesting a duplication event in the lineage leading to appendicularians.

**Figure 3.**
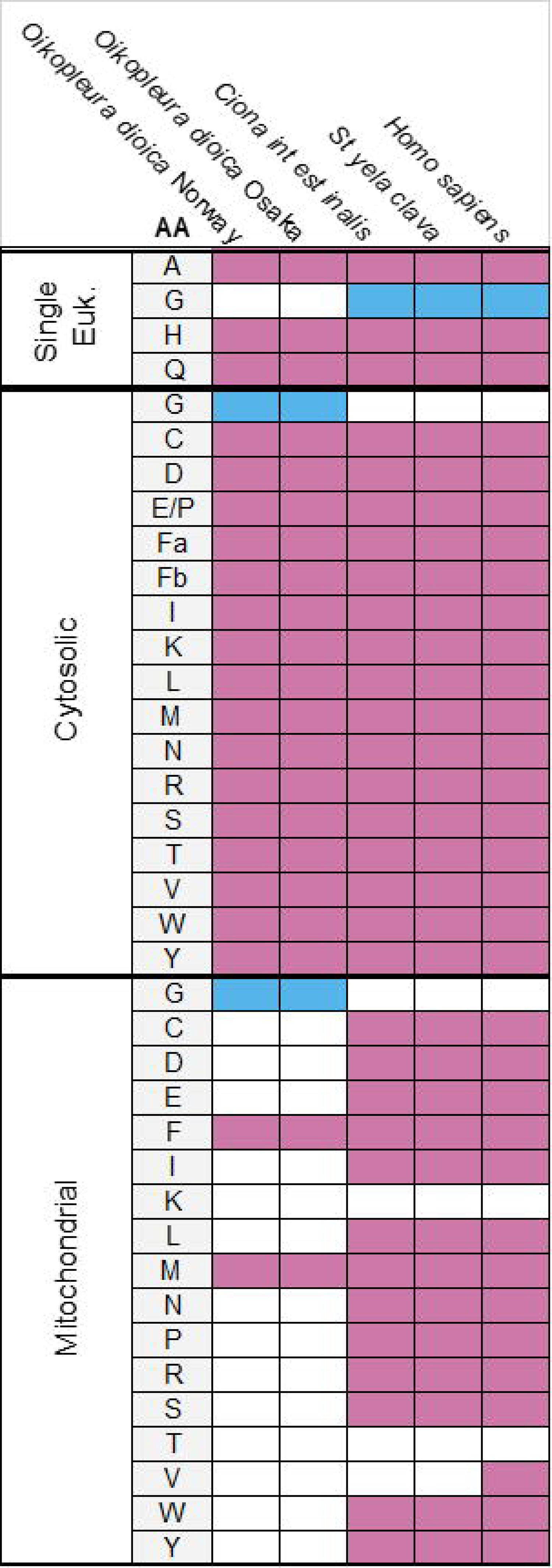
Presence-absence of aaRS orthologs in different species. Single eukaryotic (Euk): single copy aaRSs that function both in the cytosol and mitochondria. Other aaRSs are either expressed only in the cytosol (cytosolic) or in the mitochondrion (mitochondrial). aaRSs presence is shown in pink, aaRSs absence in white, and aaRS-Gly in blue. Protein accession numbers are provided in Supplementary table S5, Supplementary Material online.

### Gene Transfer to the Mitochondrial Genome

HMMs were used to search for the missing canonical mitochondrially-encoded genes (i.e., *nad2*, *nad3*, *nad4L*, *nad6,* and *atp8*) in the nuclear genome. Using these models, we found a protein sequence with a significant similarity (E-value < 5e-07) to the NAD3 model in the two published proteomes of *O. dioica* (proteins CAG5094752 and CBY12996 for the Norwegian and Okinawan genomes, respectively). Domain searches, using Cdsearch (Wang et al. 2022), confirmed that the protein sequences identified include a NAD-ubiquinone/plastoquinone oxidoreductase, chain 3 domain. These searches were conducted using the CAG5094752 and CBY12996 protein sequences as queries against the CDD v3.21 database and yielded E-values of 4.94e-03 and 3.80e-03, respectively. This protein was predicted to include two putative transmembrane domains, in support of its role in the electron transport chain (supplementary Fig. S7, Supplementary Material online). This protein-coding gene was found to be encoded on the X specific region of the X chromosome of *O. dioica* and to harbor three introns. Conversely, no nuclear-encoded protein showing significant similarity to the NAD3 model was found in the proteomes of *C. intestinalis*, *S. clava*, and *H. sapiens*.

To confirm that the identified NAD3 proteins of the Norwegian and Japanese *O. dioica* are expressed in the mitochondria we searched for the presence of mitochondrial target peptides. A significant target peptide signal was found in these putative NAD3 proteins, with a likelihood above 0.89. We could not confirm the transfer of other missing genes (i.e., *nad2*, *nad4L*, *nad6,* and *atp8*) to the nuclear genome, possibly due to the fast evolutionary rate of *O. dioica* and the short length of these proteins (150-152 amino-acid), or to the incompleteness of proteomic data for *O. dioica*.

### Phylogenetic Reconstructions

Phylogenetic reconstructions were performed using two datasets. In the first, *Doliolum nationalis* was the sole representative of the tunicate class Thaliacea. The second dataset differed from the first in including two additional thaliacean sequences: the fast-evolving sequences of *Salpa thompsoni* and *Salpa fusiformis* (supplementary table S4, Supplementary Material online). Phylogenetic trees were reconstructed under the maximum likelihood (ML) criterion using various models of sequence evolution implemented in IQtree (Minh et al. 2020; Wang et al. 2017), including complex models (i.e., MtZoa+F+G+I, MtZoa+EHO+F+G+I, Poisson+C10+G, Poisson+C20+G), and under the Bayesian CAT model implemented in Phylobayes (Lartillot and Philippe 2004). In all trees, the monophyly of tunicates was supported with maximal support (bootstrap percentage (BP) = 100, posterior probability (PP) = 1.0). However, the position of *O. dioica* was highly variable, depending on the dataset and the evolutionary model chosen (Figure 4, supplementary Fig. S8, Supplementary Material online). When the tree was reconstructed without members of the *Salpa* genus, *O. dioica* was the first tunicate to diverge when using the MtZoa+EHO+F+G+I or the mtZOA+F+I+G4 models (see Figure 4 and supplementary Fig. S8A, Supplementary Material online, respectively). Remarkably, the branch support values for a sister-clade position of *O. dioica* to all other tunicates were low (BP < 50%). Conversely, support for the monophyly of accepted tunicate clades (i.e., Stolidobranchia and Thaliacea+Phlebobranchia+Aplousobranchia) was moderate to high (BP = 60-93%). When using more complex models such as the Poisson+C10+G or the Poisson+C20+G (supplementary Fig. S8B-C, Supplementary Material online) or the Bayesian CAT model (supplementary Fig. S8H, Supplementary Material online), *O. dioica* branched within the Stolidobranchia. This positioning and the resulting relationships among Stolidobranchia clades were not supported (BP < 50%; PP < 0.5). When members of the genus *Salpa* were included in the phylogenetic analyses, *O. dioica* systematically grouped with the *Salpa* members except when using the most complex Poisson+C20+G ML model (supplementary Fig. S8G, Supplementary Material online). Interestingly, support for this relationship decreased with the complexity of the ML model used (BP = 85%, MtZOA+G+F; BP = 77%, mtZOA+EHO+F+I+G4, BP = 51%, Poisson+C10+G) (supplementary Fig. S8D-F, Supplementary Material online). In the tree reconstructed with the Poisson+20+G model, the Thaliacea monophyly was recovered (BP = 59%), but *O. dioica* branched within the Stolidobranchia and, the relationships among tunicate classes were not resolved (BP < 50%). When using the Bayesian CAT model, the phylogenetic position of *O. dioica* could not be resolved when *Salpa* was included. Specifically, convergence parameters suggested that convergence was achieved. However, different topologies were obtained depending on the chains considered. With chains 1, 2, and 3 the Thaliacea monophyly was recovered but not supported (PP = 0.53). When chains 1, 2, and 4 were considered, *Salpa* and *O. dioica* were found to be sister clades (PP = 0.57) (supplementary Fig. S8I-J, Supplementary Material online). The different branching positions of *O. dioica* among tunicates are summarized in supplementary Fig. S8K (Supplementary Material online).

**Figure 4.**
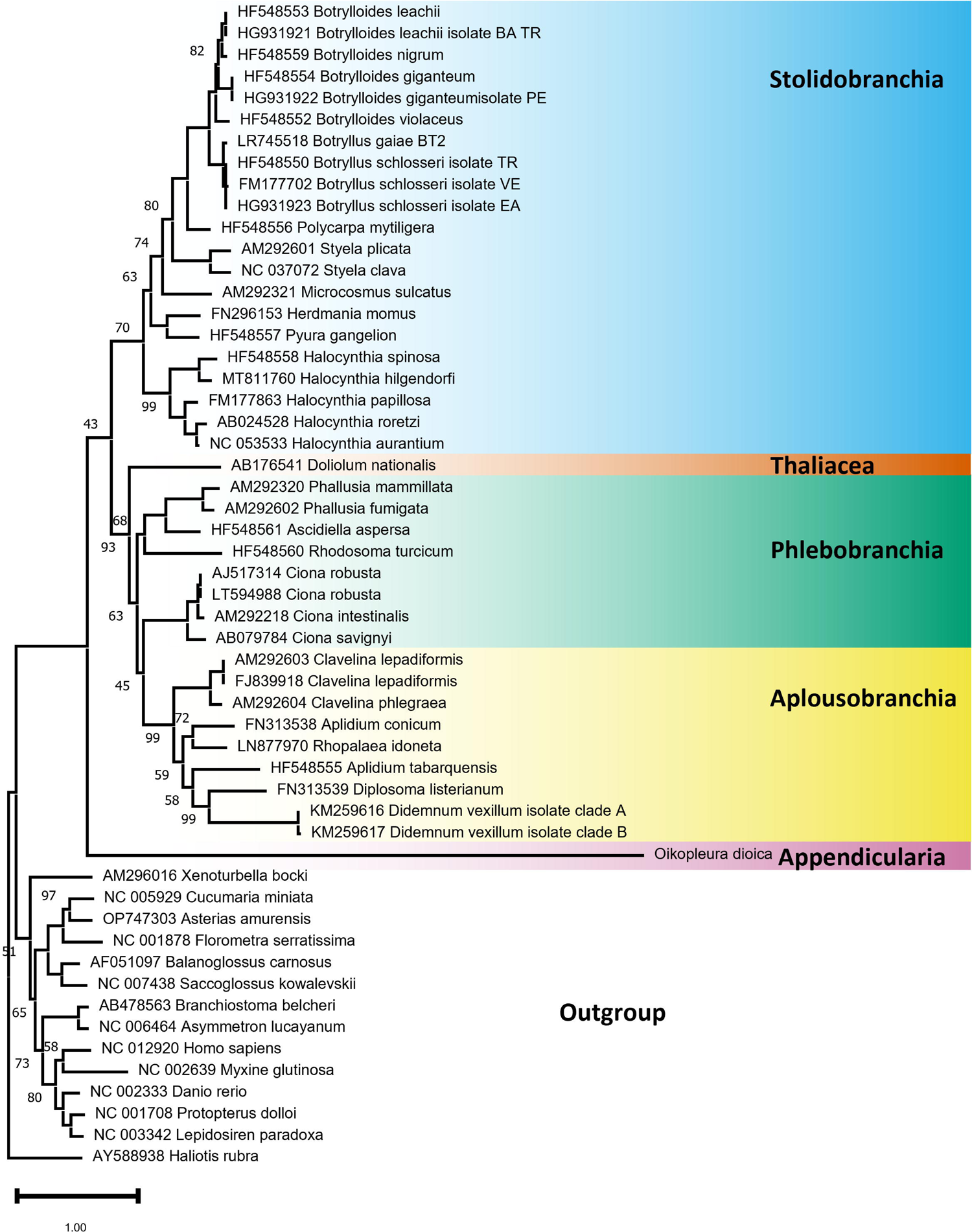
The position of *O. dioica* among chordates. The phylogenetic tree was reconstructed based on mitochondrial protein sequences under the mtZOA+EHO+F+I+G4. Branch supports are only indicated for those cases in which the bootstrap support values were lower than 100%. The tunicate classes: Stolidobranchia in blue, Thaliacea in red, Phlebobranchia in green, Aplousobranchia in yellow, and Appendicularia in purple.

## Discussion

The mitochondrial genome of *O. dioica* is extremely fast-evolving and characterized by the presence of numerous poly-A/T stretches undergoing RNA editing. Several canonical metazoan mitochondrial protein-coding genes and almost all tRNA genes are absent from the genome; there is a substantial reduction in the size of the rRNA genes; and there is evidence of a protein-coding gene transfer to the nuclear genome. Such high mitochondrial genome plasticity characterizes fast-evolving animal clades (Pett et al. 2011; Yahalomi et al. 2017).

### Protein-Coding Genes

Five canonical protein-coding genes are absent (*atp8, nad2, nad3, nad4L,* and *nad6*) from the mitochondrial genome of *O. dioica*, of which only *nad3* was identified in the nuclear genome. The four absent genes may be encoded in either the mitochondrial or the nuclear genome, but we were unable to identify them. Indeed, the *O. dioica* mitochondrial and nuclear genomes are extremely fast evolving and, consequently, these genes may have been missed by algorithms for remote homology detection, including HMM searches. Notably, these missing genes (*atp8, nad2, nad4L,* and *nad6*) are among the fastest evolving mitochondrial genes (Castellana et al. 2011; Pesole et al. 1999). The unassigned gene *ODI_mt_ORF196* is predicted to encode a protein harboring four alpha helices (supplementary figure S4, supplementary material online), unlike all other missing proteins in tunicates: the NDH2 protein of most metazoans harbors six alpha helices, while the other canonical mitochondrial proteins are predicted to harbor three or fewer alpha helices (The UniProt The UniProt Consortium 2022). Moreover, none of the methods used (Blastp, HMMer, and Cdsearch) detected a significant similarity with any other canonical proteins. We thus refrained from annotating this gene as one of the four missing genes, although Denoeud et al. (2010) annotated this gene as *nad2*.

After removing the poly-A/T regions, the largest non-coding region identified was 206 bp (between *ODI_mt_ORF196* and *cytb*) and it is possible that at least one among the *atp8* and *nad4L* genes, which are shorter than 300 bp in most tunicates, are encoded within this region. The *atp8* gene, in particular, has been overlooked in the past from the original annotation of several mitochondrial genomes (e.g. Helfenbein et al. 2004; Hoffmann et al. 1992; Le et al. 2002; Okimoto et al. 1992), including tunicates (Yokobori et al. 1999; Yokobori et al. 2003) and was then identified in ascidians following mitochondrial EST analyses (Gissi et al. 2004; Gissi and Pesole 2003).

Our results suggest that *nad3* has been relocated to the nuclear genome. While this is the first evidence of relocation of this mitochondrial gene to the nucleus in animals, a case of nuclear *nad3* has been reported in the green alga *Chlamydomonas reinhardtii* together with the transfer of *nad4L* (Cardol et al. 2006). Similar transfers to the nuclear genome have been described for other animal mitochondrial genes, for example, the *atp6* and *atp9* genes of the ctenophore *Mnemiopsis leidyi* (Pett et al. 2011) and the mitochondrial gene *atp9* in the demosponge *Amphimedon queenslandica* (Erpenbeck et al. 2006). Short ORFs are more difficult to identify than long ones, and because *atp8* and *nad4L* are relatively short it is possible that this factor, in addition to their high rate of evolution, has impaired their identification. The fact that *nad6* was detected neither in the mitochondrial genome nor in the nuclear one, is more surprising. This gene is long in other tunicates (450.9 ± 20.2 bp), and except for the gene ODI-mt-*ORF196,* no other putative ORFs longer than 300 bp were detected. The possible incompleteness of the two public nuclear genome assemblies of *O. dioica* and their inferred proteomes, or the specific criteria and bioinformatic pipeline used to annotate the protein genes in these nuclear assemblies, could also explain the failure to identify *nad6*, *atp8*, and *nad4L* in the available nuclear assemblies.

A third explanation could be that the genes that we could not identify have become fused with other mitochondrial genes. For example, the ORF we annotated as *atp6* could include regions corresponding to both *atp6* and *atp8*. An example of ORF fusion is found in the *cox*1/2 gene of amoebas (e.g., Lonergan and Gray 1996). However, this explanation is unlikely in our case since all *O. dioica* ORFs are shorter than their tunicate counterparts (Table 1), while in the case of a fusion, we would expect these ORFs to be longer.

### rRNA Genes

Tunicate rRNA genes are known to be poorly conserved. As a case in point, the boundaries of tunicate mitochondrial rRNA genes have been mostly inferred from the start and end of the flanking genes, indicating that the length estimates of rRNA genes are error-prone (Rubinstein et al. 2013; Singh et al. 2009). Only for the ascidians *Halocynthia roretzi* and *C. intestinalis*, could the rRNA boundaries be better defined, based on EST transcript data (Gissi and Pesole 2003). Although the two rRNA genes are retained in all tunicate mitochondrial genomes, they exhibit a great range of sizes. Tunicates present the shortest mitochondrial rRNA genes among chordates, with lengths varying from 596 to 984 bp for *rns* and from 921 to 1,299 bp for *rnl* (Table 1). Both rRNAs are considerably shorter in *O. dioica,* with the *rns* and *rnl* being 242 bp and 215 bp, respectively, following removal of the poly-T stretches (Table 1). Additionally, they have lost most structural similarity with other metazoan rRNA genes (HMM searches could only identify a part of the *rnl* gene). Interestingly, extreme mitochondrial rRNA reduction and lack of structural similarity are characteristics shared between appendicularians and ctenophores (Arafat et al. 2018), another lineage with an extreme rate of mitochondrial genome evolution. We annotated the two rRNA regions as different genes because most mitochondrial genomes contain these two genes. However, given the absence of sequence similarity, we cannot exclude the possibility that the *rns* region may instead be a fragment of the *rnl*. Indeed, in calcareous sponges and Placozoa, rRNA genes are fragmented and encoded on different regions of the mitochondrial genome (reviewed in Lavrov and Pett 2016). Of note, *trans*-splicing is unknown in animals.

### tRNA Genes

Tunicate tRNA gene content is variable and ranges from 22-27 tRNAs (Iannelli et al. 2007; Rubinstein et al. 2013). Specifically, the loss and acquisition of additional tRNA genes seem to be tolerated and the number of tRNA genes has been found to vary in a species-specific manner (Gissi et al. 2008; Rubinstein et al. 2013). In *O. dioica* the situation is again extreme, as all tRNAs are absent except for tRNA-Met (UAU). Nuclear-encoded cytoplasmic tRNAs import to the mitochondria has evolved many times over the course of mitochondrial evolution (Adams and Palmer 2003). We thus assume that the missing mitochondrial tRNAs in *O. dioica*’s mitochondrial genome are encoded in the nuclear genome and imported to the mitochondria.

The retention of a single tRNA-Met in the mitochondrial genome, despite the loss of all or most tRNAs, is not novel to appendicularians and has been observed in cnidarians, certain chaetognaths, and Keratosa sponges (Lavrov et al. 2023). The conventional explanation is based on the fact that mitochondrial protein synthesis starts with N-formylmethionine, as in prokaryotes, rather than methionine, the initiating amino acid in eukaryotic cytosolic protein synthesis. The addition of a formyl group to methionine is catalyzed by the enzyme methionyl-tRNA formyltransferase (Bianchetti et al. 1977). As found in prokaryotes, and in accordance with the prokaryotic endosymbiotic origin of mitochondria, this modification occurs not on free methionine but only after methionine is loaded onto a specific tRNA: namely, the initiator tRNA-fmet. Hence, it is usually assumed that the retained tRNA-Met in mitochondria is the initiator tRNA-fmet (e.g., Beagley et al. 1998).

Interestingly, animal mitochondrial genomes usually retain tRNAs that are specific to their mitochondrial genetic codes. For example, most cnidarians retain the tRNA-Trp gene necessary for the translation of the UGA codon, which is a stop codon in the nuclear genetic code (Beagley et al. 1998). In the mitochondrial genetic code of ascidians, four codons have a different assignment compared with the nuclear genetic code: methionine (AUA), tryptophan (UGA), and glycine (AGA and AGG). We thus expected the presence of a tRNA-Trp (for the translation of the UGA codon) and a tRNA-Gly (to translate the codons AGA and AGG) in the mitochondrial genome of *O. dioica*. Therefore, we computed the codon frequency across all protein-coding genes of *O. dioica*, thus verifying the usage of all these AUA, UGA, AGA, and AGG codons in the mitochondrial genome (supplementary table S6, Supplementary Material online). Indeed, it has been found that translation of the AGA and AGG codons to Gly improves the similarity between *O. dioica* and tunicate mitochondrial proteins, supporting the use of the ascidian mitochondrial genetic code in appendicularians (Pichon et al. 2020). The presence of a cytosolic and a mitochondrial aaRS-Gly, observed while searching for mitochondrial aaRS in the *O. dioica* nuclear genome, further supports the view that a specific mitochondrial tRNA recognizes the glycine AGA and AGG codons. Regarding protein-coding genes, two explanations are possible for the lack of mitochondrial Gly and Trp tRNAs: either we failed to identify these tRNA genes in the sequenced mitochondrial genome due to their very unusual secondary structure or because they are interrupted by poly-A/T stretches; or these tRNAs are nuclear-encoded and imported from the cytosol. In support of the latter scenario, it was found that among animals, both ctenophores and myxozoans have lost all their mitochondrial tRNA but still use the UGA codon as tryptophan (Arafat et al. 2018; Pett et al. 2011; Yahalomi et al. 2017).

### Genome Editing of Poly-A/T Stretches

The sequencing of PCR fragments using ONT long-reads allowed us to assemble a large part of the mitochondrial genome of *O. dioica*. It confirmed the presence of numerous poly-A/T stretches in the DNA sequence that are reduced to 6U-sites in the RNA transcript. Because accurate sequencing of long homopolymers is currently unfeasible using Sanger, Illumina (Ari and Arikan 2016), or Nanopore sequencing (Delahaye and Nicolas 2021; Sereika et al. 2022), assessing the length of T/A stretches in our assembly is unreliable. The average size of these regions is likely slightly higher than that reported in the assembly because sequencing methods tend to underestimate the length of homopolymer stretches. Another source of uncertainty regarding the length of the homopolymer stretches stems from the fact that we sequenced amplicons obtained from DNA extracted from a pool of polymorphic individuals. Intra-species variation in the length of these stretches likely exists, further contributing to inference uncertainty.

What is the molecular mechanism underlying the editing to 6U-sites and, more specifically, how are these sites recognized? We could not identify a conserved sequence motif surrounding these sites (Supplementary Fig. S3, Supplementary Material online). Our analysis revealed the following general motif for the poly-A/T stretch (see Materials and Methods): (1) the 5’-end of this strand starts with a perfect 6T homopolymer; (2) downstream of this region, there is a poly-pyrimidine track whose length varies from 2 to 509 bases; (3) it ends when a purine is first encountered. Only the poly-pyrimidine track is excised. We speculate that the perfect 6T motif in the upstream region may be the beginning of the recognition editing signal and that a purine marks its end. In between these motifs, the polypyrimidine track is excised. Notably, the enzymes involved in this excision mechanism are still unknown. Another kind of editing was observed within the cytochrome *b* gene: rather than a perfect 6T motif, it starts with “TTTTTCTT”, followed by an irregular excised region, “AACTT”, as it contains two purines (supplementary Fig. S9, Supplementary Material online).

Mitochondrial genome editing is rare in animals and is usually limited to the insertion of a few bases (Lavrov et al. 2016). Here, rather, the deletion of a large polypyrimidine region is inferred. The removal of such a long stretch at the RNA level is a characteristic of introns (Chorev and Carmel 2012). Spliceosomal introns are characterized by short signal sequences such as GT/AG and AT/AC (Berná and Alvarez-Valin 2014; Rogozin et al. 2012). The RNA editing, we observed here, seems to differ greatly from such a splicing mechanism, as we could not find similar motifs. Similarly, the poly-A/T stretches bore no sequence similarity to self-splicing introns, which have well-defined secondary structures (Waring and Davies 1984).

Trypanosomatid protists present a cleavage-ligation pathway through which multiple uridines can be deleted or inserted by a U-specific exonuclease and a terminal uridil transferase, respectively (Cruz-Reyes and Sollner-Webb 1996). This particular editing mechanism involves guide RNAs, which determine the number and positions of the U nucleotides deleted or inserted within the mitochondrial genome of these protists, e.g., *Leishmania tarentolae* (Connell et al. 1997). Future studies are necessary to determine whether some components of the editing mechanisms in these protists and in *O. dioica* are evolutionarily related.

Appendicularians have classically been divided into three families: Fritillariidae, Kowaleskiidae, and Oikopleuridae (Garić 2024). However, preliminary results from our group suggest that the Kowaleskiidae are actually members of the Fritillariidae (Garić et al. 2018). The presence of edited homopolymer stretches has been noted for the COI gene of *O. vanhoeffeni* (Latour 2021). Interestingly, we observed that large poly-A/T inserts are also present in members of the Fritillariidae, suggesting that this mitochondrial editing mechanism was present in the ancestor of Appendicularia (Garić et al. 2018). Conversely, while examining published data from Naville et al. (2019), we identified a clade within the Oikopleuridae that includes the species *Bathochordaeus stygius*, *Mesochordaeus erythrocephalus*, and *Oikopleura longicauda.* These three species do not harbor large poly-A/T stretches (Garić et al. 2018), suggesting that this RNA editing mechanism may have been lost in Oikopleuridae. Additional genomic and transcriptomic data from a larger taxon sampling of appendicularians are needed to better understand the distribution and evolutionary dynamics of this editing mechanism.

EST and transcriptomic RNAseq data are available for *O. dioica* from Norway and mainland Japan, respectively. For the Norway sample, the genomic data obtained in this work indicated the presence of RNA editing. To evaluate whether the poly-A/T stretches are conserved among *O. dioica* lineages, we compared the location of 6U-regions in the DNA and transcriptomic data from Norway and mainland Japan, respectively (Supplementary Fig. S2, Supplementary Material online). Our results suggest that while many of the 6U-loci are conserved, differences do exist. For example, in the *cox1* gene, a 6U-site in the Norwegian sequence and a UUCUUU site in the Japanese sequence can be reliably aligned. These differences suggest that the poly-A/T stretches loci may not be conserved in the mitochondrial genomes of different populations or species. Of note, it was recently suggested that *O. dioica* from Japan and Norway represent different species (Masunaga et al. 2022; Plessy et al. 2024). The birth and death dynamics of these poly-A/T stretches will be better clarified when genomic and transcriptomic data from a larger appendicularian taxonomic sampling become available.

That we were able to assemble large contigs encompassing several genes from RNA sequences indicates that the pre-mRNA transcripts in *O. dioica* are polycistronic. In vertebrates, it has been shown that the cleavage of tRNAs that flank protein-coding genes constitutes the mechanism (termed tRNA punctuation) that releases monocistronic or bicistronic mature mRNAs of mitochondrial coding genes from the large RNA precursor (Ojala et al. 1981). In ascidians, the transcription and RNA maturation process of the mitochondrial genome has been reconstructed through EST analyses, revealing that the main features of mitochondrial transcription are conserved between vertebrates and ascidians (*i.e.,* the presence of polycistronic and immature transcripts; the creation of stop codons by polyadenylation; tRNA signal punctuation; and rRNA transcript termination signals) (Gissi and Pesole 2003). Since most tRNAs have been lost in *O. dioica,* a mechanism differing from the tRNA signal punctuation is likely employed to release mature mRNAs.

### The Phylogenetic Position of *O. dioica*

Based on both molecular (Delsuc et al. 2018) and morphological (Braun et al. 2020) analyses, *O. dioica* was placed as a sister-clade to all other tunicates. However, it was recognized that this placement, in molecular studies, may stem from a long-branch-attraction (LBA) artifact (Delsuc et al. 2018). In our analysis, we found that the phylogenetic position of *O. dioica* was highly variable (supplementary Fig. S8K, Supplementary Material online). The grouping of *O. dioica* and members of the genus *Salpa* is most likely an LBA artifact (Philippe et al. 2005). However, even when the two *Salpa* representatives were removed from the phylogenetic analyses, the phylogenetic position of *O. dioica* was never well supported and depended on assumed probabilistic model used. The lack of a phylogenetic signal in the mitochondrial dataset, which precludes the reconstruction of phylogenetic relationships at the phylum level, is a feature of fast-evolving species (Arafat et al. 2018; Pett et al. 2011; Yahalomi et al. 2017).

### An *O. dioica* Mitochondrial Genome from Okinawa, Japan

After completing this manuscript, a description of the complete mitochondrial genome of *O. dioica* from Okinawa (Japan) (accession number PP146516) was reported in a preprint (Dierckxsens et al. 2024). While numerous substitutions differentiate the two genomic sequences (113 bp difference in the barcoding region corresponding to a K2P distance of 20.3%), the gene order was identical. One difference is that of the presence of a region containing several tandem repeats between the *cox1* and the gene we have labeled ODI_mt_ORF196. Remarkably, Dierckxsens et al. (2024) succeeded in assembling a circular molecule. The region in the Okinawa sample that is homologous to the part that we failed to sequence, is a large non-coding region of ∼28,000 bp composed of numerous repetitive elements. A similar structure may be present in the Norway sequence analyzed here.

## Conclusions

We have demonstrated that a combination of DNA and RNA data from long- and short-read sequencing could significantly improve our understanding of the structure and gene content of the *O. dioica* mitochondrial genome.

The mitochondrial genome of *O. dioica* shows a unique gene content and an extreme rate of sequence evolution. A total of 28 genes are missing, including *atp8*, *nad2*, *nad3*, *nad4L*, *nad6*, and all tRNA genes except tRNA-Met (UAU). The mitochondrial gene *nad3* was transferred to the nucleus and harbors a mitochondrial target peptide. Further work is however needed in order to decipher the complete sequence of the mitochondrial genome, and in particular to unravel the accurate sequence and length of the poly-A/T regions. Unfortunately, no sequencing techniques currently unable us to perfectly sequence these regions, as the sequencing methods Illumina, ONT, and Sanger approaches failed to correctly sequence long polynucleotide regions. However, information obtained from the sequencing of PCR fragments helped us to assemble a draft mitochondrial genome.

Because of the fast-evolutionary rate, we were unable to accurately determine the phylogenetic position of *O. dioica*. LBA artefacts are expected due to such a high evolutionary rate, and the inferred tree often varied depending on the set of species included in the analysis. Thus, mitochondrial sequences are not the markers of choice when the goal is the accurate inference of phylogenetic relationships within appendicularians. We expect that the availability of nuclear genomes whose rate of evolution is lower than that of the mitochondrial genome, will enable more accurate reconstruction of the phylogenetic relationships among appendicularian species. Of note, comparative genomic studies of *O. dioica* and additional members of the appendicularians are expected to provide us with a better understanding of the evolution of RNA editing in this group.

## Materials and Methods

### DNA Extraction, Sequencing, and Assembly

Living specimens of *O. dioica* were provided by the Sars International Centre for Marine Molecular Biology (SICMMB) (Bergen, Norway) in March 2019 and cultured at Humboldt-University zu Berlin (HU) following the methods detailed in Bouquet et al. (2009). The animals sequenced in this study originated from a cross between new animals from the field collected at Rosslandspollen, north of Bergen (60°33’54.6“N 5°2’24.5”E), and an existing culture (generation 29). These animals were grown for ∼20 generations in the lab. Ethanol-preserved samples were shipped to Tel-Aviv University. Genomic DNA was extracted from ∼100 whole individuals following the protocol of Fulton (2012). About 100 ng were sent to the Technion sequencing center (Haifa) for Illumina sequencing (November 2019), the libraries were prepared using the TruSeq DNA Nano library prep kit (Illumina), and were sequenced on an Illumina Hiseq 2500 platform. Similarly, about 1 μg of total DNA was sent to Hy Laboratories Ltd (Hylabs, Rehovot) for ONT sequencing using a MinION Mk 1 R9.4 flow cell with a MinION Mk1B device and 1D sequencing kits (May 2019). However, the ONT sequencing only provided 962 reads (mean length 750 bp). ONT sequencing was thus performed again using a different DNA extract and a different sequencing service (the Technion sequencing center) (on July 2019 and December 2021). The flow cell and device setting and sequencing kits were the same as previously. The first sequencing (July 2019) provided ∼2 million reads (mean length 2,330 bp) but we could not assemble the mitochondrial genome using these data. *A posteriori,* we found that only ∼600 reads mapped to the mitochondrial genome (mean length 300 bp) and that these reads only mapped to a few regions of the mitochondrial genome (data not shown). For this second ONT sequencing, fragments around 3,000 bp were selected. Because we realized that mitochondrial DNA might be fragmented, we performed the last ONT sequencing (from December 2021) without fragment size selection. This ONT sequencing also provided only 3,144 reads (mean length 1,070 bp).

The Illumina 100 bp paired-ended reads were cleaned from adaptors using Cutadapt v.2.8 (Martin 2011) and assembled using IDBA-UD v.1.1.3 (Zhu et al. 2010) with default parameters. Efforts to assemble the mitochondrial genome from these reads were unsuccessful because contigs were broken at the long poly-A/T regions.

Neither Illumina nor ONT sequencing enabled us to assemble a complete mitochondrial sequence. We thus decided to amplify the mitochondrial genome from several PCR fragments. To do so, we needed to design primers in regions that did not include homopolymers. We used published RNA reads and EST sequences to identify mitochondrial genes and design primers.

### Identification of Canonical Mitochondrial Protein Transcripts in Published RNA and EST Sequences

Illumina RNA reads from *O. dioica* specimens from Japan and the North Sea were downloaded from the National Center for Biotechnology Information (NCBI) (accessions: SRR1693762, SRR1693765, SRR1693766, and SRR1693767) (Wang et al. 2015), and the European Nucleotide Archive (ENA) (accessions: SRR20015061) (Raghavan et al. 2023), respectively. These reads were assembled using IDBA-UD v.1.1.3 (Zhu et al. 2010) with default parameters. tBlastn searches were conducted against the two transcriptome assemblies obtained using the amino acid sequences of the protein-coding genes of the mitochondrial genome of *Ciona robusta* (AJ517314) as query. The reads from Japan could be assembled in four contigs and will be referred to as “Japan” (supplementary Fig. S10A, Supplementary Material online). The genes from the North Sea could be assembled in five contigs and will be referred to as “North Sea” (supplementary Fig. S10B, Supplementary Material online). All contigs were annotated with the webserver MITOS V2 (Donath et al. 2019).

Mapping Illumina DNA reads and the few ONT reads to these contigs confirmed the presence of homopolymers and also indicated that the Japanese *O. dioica* presented many substitutions when compared to our sample, which originated from Norway. EST sequences from *O. dioica* samples from Norway were available in the OikoBase database (Danks et al. 2012). These data originated from natural populations of *O. dioica* caught in the fjords near Bergen, Norway (Denoeud et al. 2010). For each mitochondrial ORF identified in the Illumina North Sea dataset, we mapped ∼500 EST sequences to the reference. We next filtered the flanking regions of the extracted EST sequences against the UniVec Vector database using Geneious Prime v.2023.2.1. The consensus sequences were extracted for each gene, which helped us determine the exact gene boundaries. We verified that all coding transcripts we had identified could be successfully translated to proteins using the ascidian or the invertebrate mitochondrial genetic codes. We found that the ascidian genetic code yielded proteins with a higher sequence similarity to mitochondrial proteins in other animals than when using the invertebrate genetic code, in agreement with Pichon et al. (2020).

### PCR Amplification, Sequencing, and Assembly

Primers of ∼ 30 bp with a TM ∼ 60 C° were designed based on the ORFs assembled from the EST, accounting for the gene order inferred from the contigs (supplementary table S1, Supplementary Material online). Long PCRs were performed with several primer combinations (one from each pair of contigs and in various orientations), to determine gene arrangements along the mitochondrial genome. Because of the high AT content of the genome, we were unable to design primers that would result in overlapping PCR fragments. Instead, we used the same region for forward and reverse primers, leading to non-overlapping fragments. The sequence of the primer region was confirmed using Illumina data. PCRs were performed in a 25μl volume with 2u of TaKaRa Ex-Taq polymerase (Cat.# RR001), 1X of Ex-Taq buffer, 1M of Betain, 0.2mM of dNTPs (TaKaRa mixture), 0.5μM of each primer and 1μl *O. dioica* DNA extract. The PCR program used: (a) a long denaturation step at 95°C for 5 min; (b) 35 cycles of a short denaturation step at 95°C for 45 sec followed by an annealing plus elongation step at 68°C for 10 min; and (c) a long elongation step at 72°C for 10 min. Reamplifications were performed to improve the PCR yield of some fragments. Reamplifications were performed using 1 μl of PCR products in place of genomic DNA. Twelve PCR reactions were successful and sent for ONT sequencing (supplementary Fig. S1, Supplementary Material online). Each PCR product was barcoded and the sequencing was performed on May 2023 by the Technion Sequencing Center on an Oxford Nanopore MinION device (MinKNOW software v.22.12.7), using one R9.4.1 flow cell (Oxford Nanopore, cat no. FLO-MIN106D). The ONT reads obtained were filtered from adaptors using Porechop v.0.2.4 (Wick et al. 2017) with the parameter - middle_threshold 85. Next, we assembled the trimmed ONT reads with Flye v.2.9.1 (Kolmogorov et al. 2020) as implemented in Geneious Prime v.2023.2.1 with two rounds of polishing. The same parameters could not be used for all PCR fragments, as in some cases the assembly program returned an error message or failed to terminate. In these cases, other parameters were tested: e.g., if the default parameter “genome size” of 1,000 bp resulted in a failed run, we tried running the program with increasing values of genome size from 2,000 to 5,000 bp. If this also failed, we added the option “metagenome” or the option “reduced contig = 30”. However, even with these modifications the assembly of the PCR fragment #12 was unsuccessful.

Many of the assembled contigs were palindromic, with the middle of the contig corresponding to the PCR primer. In this case, the consensus sequence was split into two halves, reads were mapped to each half (using Geneious Prime v.2023.2.1 with default options), and the region with the highest number of mapped reads was kept as the consensus sequence.

The entire mitochondrial sequence was determined by connecting the obtained contigs in their overlapping primer regions, i.e., the same primer region was the forward primer of one PCR fragment and the reverse primer of another fragment. The exact sequences of the primer regions were verified by mapping the Illumina reads to the assembled genome sequence.

### Annotation of the Mitochondrial Genome

#### Protein-coding genes

Gene annotation was based on Blast searches against the NCBI database and was confirmed using MITOS v.2 (Donath et al. 2019). The end positions of protein-coding genes were verified using EST data (the presence of a poly-A region after the stop codon). The start codon was determined by searching for the first start codon upstream of the region that corresponded to the EST data.

#### rRNAs

The mapping of Illumina RNA reads from Japan (SRR1693762, SRR1693765, SRR1693766, SRR1693767) and the North Sea (SRR20015061) to the mitochondrial genome of *O. dioica* revealed two regions with higher coverage that we inferred to be *rns* and *rnl* (supplementary Fig. S5, Supplementary Material online). Read mapping was done using the Geneious Mapper, Geneious Prime v.2023.2.1 with the following parameters: medium-low sensitivity, randomly map multiple best matches, only map paired reads which both map, and trim paired read overhangs. Average coverage was computed with Geneious Prime v.2023.2.1 (i.e., panel statistics). The region that we annotated as *rnl* was also identified as such using gene prediction with MITOS v.2 (Donath et al. 2019). The *rnl* and *rns* sequences were also present in the EST data, which helped determine the boundaries of these ribosomal genes.

#### tRNAs

Putative tRNAs were searched using MITOS v.2 (Donath et al. 2019) and the ARWEN v.1.2 webserver (Laslett and Canbäck 2008). Both programs identify a single tRNA gene (supplementary Fig. S6, Supplementary Material online).

### Aminoacyl-tRNA Synthetase Profiling

To corroborate the absence of tRNA genes in the mitochondrial genome, we searched for aaRS genes in the nuclear proteome of *O. dioica* (i.e., the amino acid sequences of *O. dioica* in the NCBI “protein” database). The aaRS search was done using HMMER v.3.3.2 software (Eddy 2011) and was based on profiles developed by Pett and Lavrov (2015). These profiles were used to search against proteome datasets downloaded from NCBI on April 2023 of *O. dioica* from Norway (GCA_000209535), *O. dioica* from Okinawa (GCA_907165135), *H. sapiens* (GCF_000001405.40 Human Build 38 patch release 14), *C. intestinalis* (GCF_000224145.3), and *S. clava* (GCF_013122585.1).

### Nuclear Gene Transfer

We used HMM searches against the proteome of *O. dioica* from Norway GCA_000209535 and *O. dioica* from Okinawa GCA_907165135 to detect canonical mitochondrial-encoded genes that could have been transferred to the nuclear genomes. The searches were performed with HMMER v3.4 (Eddy 2011) using the *atp8*, *nad2*, *nad3*, *nad4L*, and *nad6* gene models developed by Gabriel Alves Vieira and available in the mitos2_wrapper source code repository (Vieira 2023). As negative control we also searched the proteomes of *H. sapiens* (GCF_000001405.40 Human Build 38 patch release 14), *C. intestinalis* (GCF_000224145.3), and *S. clava* (GCF_013122585.1). Putative mitochondrial proteins that were found to be encoded in the nuclear genome were searched for mitochondrial targeting signal using the TargetP-2.0 webserver (Armenteros et al. 2019).

### Codon Usage and Amino-Acid Composition

Codon usage and amino acid composition were computed using Geneious Prime v.2023.2.1.

### Characterization of ODI_mt_ORF196

The number of transmembrane domains of ODI_mt_ORF196 was predicted using TMHMM v.1.0.2 in Geneious Prime v.2023.2.1 (Viklund and Elofsson 2004). The protein structure was predicted using Alphafold (Jumper et al. 2021). The predicted structure was used as query against a large database of structures using Foldseek (van Kempen et al. 2024).

### Characterization of the Poly-A/T Stretches

We defined poly-T stretches by manually examining 59 cases of edited poly-T sites, i.e., these editing cases were validated by comparing RNA and DNA sequences of these loci. A classic poly-T stretch is defined as follows: (1) the 5’-end of this stretch starts with a perfect 6T homopolymer. This region is not excised; (2) downstream of this region, there is a C/T stretch (i.e., a Polypyrimidine track). This region is excised; (3) the stretch ends when either an A or a G nucleotide is encountered. This A or G nucleotide is not part of the poly-T stretch and is not excised. In six validated cases this pattern was violated. In five of these cases there was a single A nucleotide within the long C/T stretch and in one case a G. We thus include in the definition also cases in which the C/T stretch comprises a single purine, as long as it is followed by at least 10 more C/T nucleotides. Searching the genome with this definition, we found six additional poly-T stretches (four classic without purine in the C/T stretch and two non-classic with one purine in the C/T stretch) for which we lack RNA support. In total, we annotated 65 poly-T stretches in the mitochondrial genome of *O. dioica*. As we searched for the above motif or its reverse complement to account for both strands, we termed these stretches poly-A/T stretches.

For each homopolymer site in the genome we extracted 50 bp before the site, the 6 T/A in the site itself, and 50 bp after the site using our own python v.3.11.5 script on Jupiter Notebook v.7.1.2. Nucleotide frequencies before and after the homopolymer sites were plotted (Supplementary Fig. S3, Supplementary Material online) using WebLogo v.2.8.2 with parameter “Frequency Plot” selected in the section “Advanced Logo Options”. (Crooks et al. 2004). To determine the conservation of the poly-A/T stretches, we aligned the mitochondrial gene regions extracted from the Illumina RNA data from Japan to our own assembly. Pairwise alignments were computed using MAFFT v.7.490 under the L-INS-i algorithm.

### Barcoding Analysis to Confirm Species Identity

We used a barcoding approach to confirm that our DNA samples and the EST deposited in Oikobase belong to the same *Oikopleura* species. We computed the evolutionary Kimura two parameters (K2P) distance using MEGA v.11.0.13 (Tamura et al. 2021) between the barcoding region of the mitochondrial sequence we had assembled (i.e., the CDS region without the poly-A/T stretches) and the consensus barcoding region we had assembled from the EST sequence. We extracted the extended tunicate barcoding *cox1* region of Salonna et al. (2021) from the mitochondrial DNA sequence. We next searched for sequence similarity against the EST data using Blastn. Out of the 100 best hits, a consensus sequence of the EST *cox1* region was computed. Specifically, the K2P distance between the two sequences (i.e., the EST consensus and the mitochondrial genome) was 0.0093 substitutions per site (6 bp differences, 0.9%). Because the number of bp differences among the EST sequences ranged from 0 to 20 (average ∼8bp), we inferred that our sample and the EST data had originated from the same species. Moreover, the barcoding region from the North Sea RNA contigs was also found to be identical to the EST consensus sequence, suggesting that these RNA reads had originated from the same species as well.

In contrast, the barcoding region of the Japanese RNA contigs presented 93 bp differences from the EST and the mitochondrial genome from Norway assembled in this work (K2P distance of above 0.163 substitutions per site). This suggests that these mainland Japanese sequences originated from a different species than the *Oikopleura* species whose mitochondrial genome was determined in this work (Masunaga et al. 2022; Plessy et al. 2024).

The sequence PP146516 from Okinawa (Dierckxsens et al. 2024) not only greatly differed from the Norway sequences, (above 111 bp difference in the barcoding region and K2P distance of above 0.198 substitutions per site) but also greatly differed from the Japanese contig we had assembled from mainland Japan (94 bp difference in the barcoding region and K2P distance of 0.163 substitutions per site), supporting the view that PP146516 belongs to a third *Oikopleura* lineage (Masunaga et al. 2022; Plessy et al. 2024).

### Mitochondrial Genome Length among Tunicates

The average mitochondrial genome size of tunicates and the average gene lengths were calculated based on 34 complete mitochondrial genomes of tunicates downloaded from the NCBI database (supplementary table S4, Supplementary Material online).

### Phylogenetic Analysis of Mitochondrial Protein Sequences

The phylogenetic analyses encompassed 34 tunicate species with a complete mitochondrial genome that were deposited in NCBI (last accessed: June 2023) and 14 outgroup species (supplementary table S4, Supplementary Material online). The outgroup species were chosen following Singh et al. (2009). Eight protein-coding gene (*atp6, cox1, cox2, cox3, cytb, nad1, nad4, nad5*) were extracted from the Genbank flatfile of each species using Geneious Prime v.2023.2.1. We used the Guidance2 server version 2.02 (Sela et al. 2015) with default parameters to identify unreliable regions of the protein alignment, which was generated using MAFFT version 7.407, also with default parameters. We excluded these regions as well as alignment columns that included more than 50% missing data or gaps. The different protein-coding alignments were then concatenated to form a super-matrix with 2,027 columns. Because members of the tunicate genus *Salpa* are extremely fast evolving, we prepared a second dataset, repeating the above steps, but excluding *S. thompsoni* and *S, fusiformi.* The resulting super-matrix of 46 species included 2,176 columns.

The phylogenetic trees were reconstructed under the maximum likelihood (ML) criterion using IQ-TREE 2.1.2 (Minh et al. 2020). First, we identified the best-fitting model by using the ModelFinder-Plus (MFP) option of IQ-TREE 2.1.2 and computed the ML tree using this model. Using IQtree, the model that best fitted both datasets (with and without the genus *Salpa*) was mtZOA+F+G+I. We also reconstructed the evolutionary relationships using complex models that are not implemented in MFP: mtZOA+EHO+F+G+I, Poisson+C10+G, and Poisson+C20+G. The robustness of each internal branch of the ML trees obtained was evaluated with 100 bootstrap replications (Felsenstein 1985).

Bayesian tree reconstructions were performed using the site-heterogeneous CAT model implemented in Phylobayes MPI v1.5 (Lartillot et al. 2013), which is expected to lower the impact of long-branch attraction on the inferred phylogenetic tree (Lartillot et al. 2007).

For the first and second datasets, four independent chains were run for 71,880 and 188,000 cycles, respectively. The first 5,000 and 10,000 trees from each chain, for the first and second datasets, respectively, were discarded as burn-in and trees were sampled every 10 trees (i.e., the sample size was 3,160, and 17,800 trees per chain for the first and second dataset respectively). Chain convergence was assessed using the bpcomp and tracecomp scripts, which are part of Phylobayes. Specifically, the bpcomp maxdiff values were 0.23 and the tracecomp effsize and rel_diff values were >800 and below 0.15, supporting a proper convergence for the first dataset. For the second dataset, the bpcomp maxdiff value was 0.40 when the four chains were considered together, which indicated insufficient convergence. However, when considering only chains 1, 2, and 3 or chains 1, 2, and 4 this value dropped to 0.18 and 0.27, indicating proper convergence. In both cases, rel_diff values were >150 and below 0.16, supporting convergence.

## Supplementary Material

Supplementary files Tables S1-S6 and Figures S1-S10 are available at Genome Biology and Evolution online.

## Acknowledgments

We want to thank Nitzan Fourier from the Technion Genome Center for her help with Nanopore sequencing. We thank Yair Shimony for his help with the structure predictions.

## Author Contributions

D.H., C.G., and T.S. conceived the study and designed the research. T.S. provided the biological samples. M.N. and Y.K. performed the bench work. D.H. and Y.K. assembled and annotated the mt genome. D.H., M.N., T.P., and Y.K. performed the various evolutionary analyses and drew the figures. All authors contributed to interpretation of the results. D.H., T.P., and Y.K. wrote the first draft of the manuscript. All authors assisted in revising the manuscript and approved the final version of the text.

## Data Availability

The Illumina read data have been deposited in the European Nucleotide Archive under study ID number PRJEB72855. The mtDNA sequence was deposited under accession number OZ035851. The ONT reads obtained for each of the PCR fragments, the phylogenetic trees, as well as the sequences assembled from published RNA and EST data are available on GitHub (https://github.com/dorohuchon/Oikopleura_dioica_mitochondrial_genome).

