## Supplementary Material online for "Evolutionary Insights from the Mitochondrial Genome of *Oikopleura dioica*: Sequencing Challenges, RNA Editing, Gene Transfers to the Nucleus, and tRNA Loss"

**This file includes:**

**Supplementary Figures S1 to S10**

**Supplementary Tables S1 to S6**

Figure S1: PCR primers and fragments

PCR fragments and primers position in *O. dioica* putative circular mitochondrial genome. Protein-coding genes are in green, rRNA genes are in dark blue, the tRNA gene is in light blue, PCR fragments are in pink, and primers are represented by purple arrows. A non-sequenced region indicated as a “gap” is in gray. The genes are pointing in the direction of their transcriptional orientation.

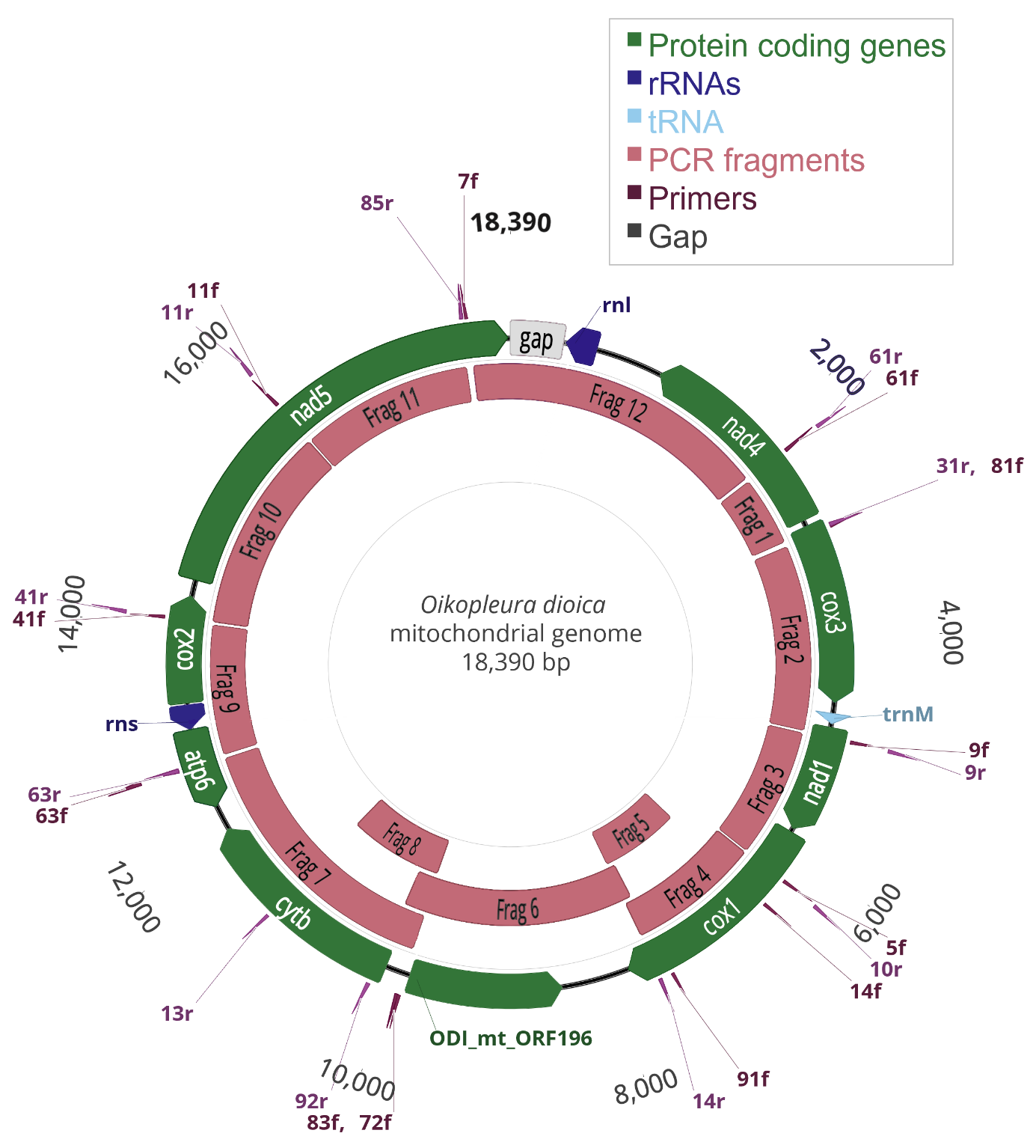

Figure S2: Comparison of poly-T versus 6U-sites location between *O. dioica* from Norway and Japan

Comparing the location of DNA poly-T and RNA 6U loci along protein-coding genes between *O. dioica* specimens from Norway and Japan, respectively. Each pair is a comparison between the DNA assembly obtained in this study from the Norwegian samples, and coding sequences obtained from RNA sequences from Japan (SRR1693762, SRR1693765, SRR1693766, SRR1693767). Light-blue arrows indicate poly-T and 6U sites.

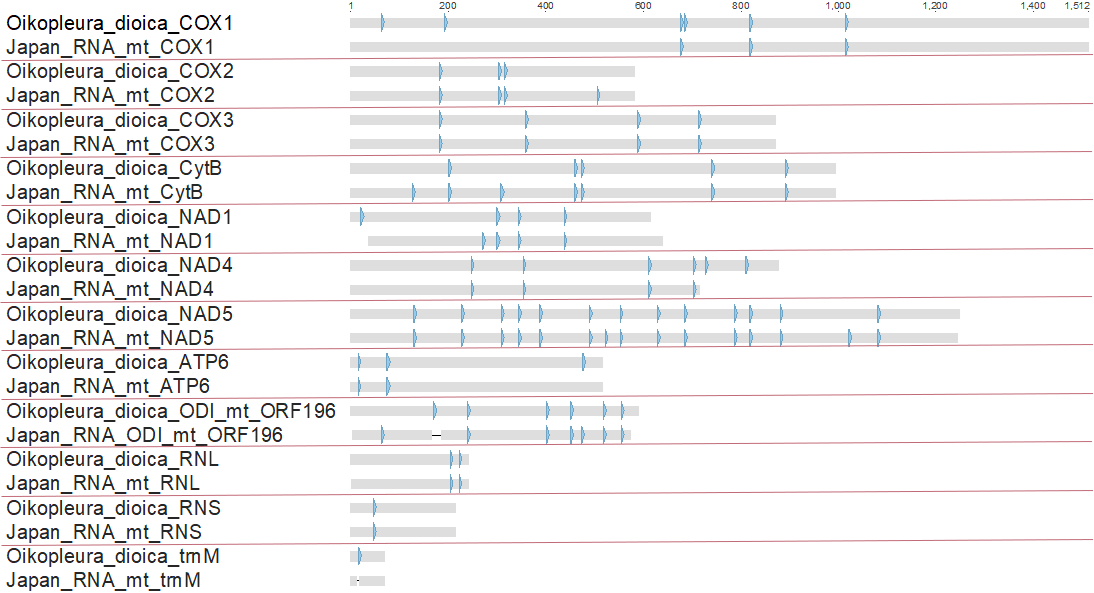

Figure S3: Nucleotide frequencies around poly-T stretches

A poly-T stretch always starts with 6 consecutive T residues followed by a tail of C/T residues with a variable length. We searched for signal before (A) or after (B) the poly T stretch (the poly-T stretch itself is now shown, and is found downstream of position 50 in (A) and upstream of position 1 in (B)). The frequency is represented by the size and height of the nucleotide logo. The frequencies are averaged over 65 poly-T regions.

**A.** Before poly-T site

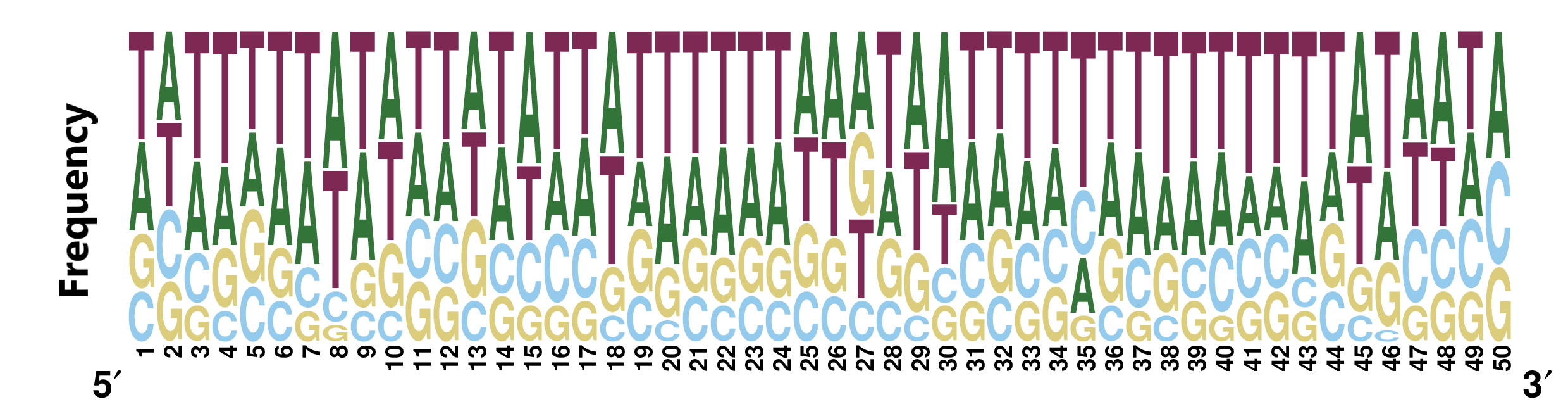

**B.** After poly-T site

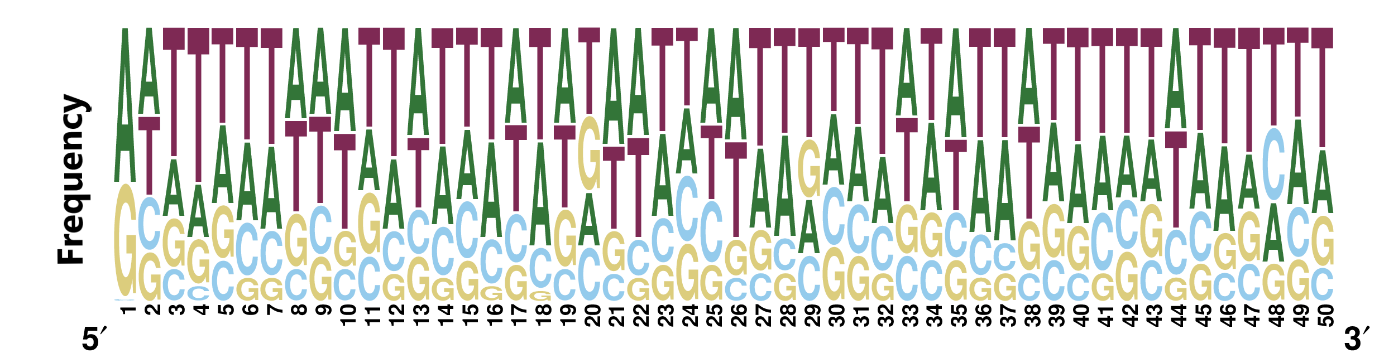

Figure S4: ODI_mt_ORF196 predicted secondary structure

ODI_mt_ORF196 protein structure predicted using Alphafold.

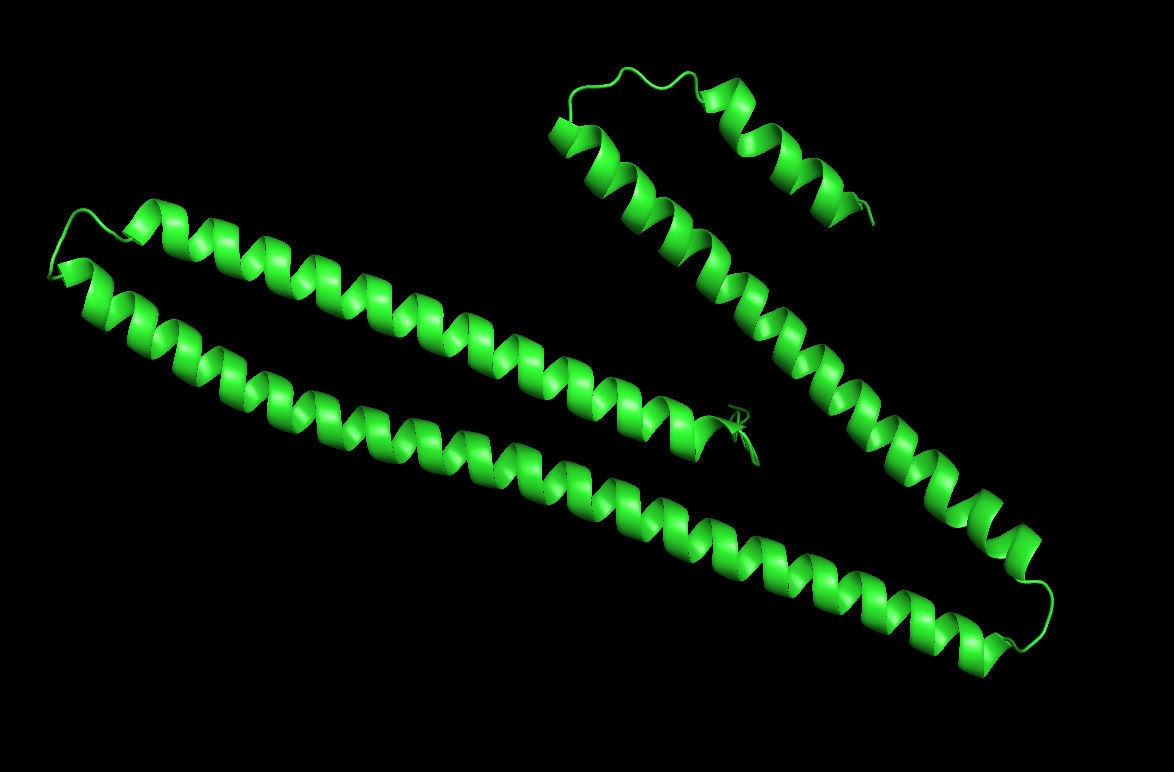

Figure S5: rRNA read coverage

Coverage (in light blue) of RNA Illumina-reads mapped to *O. dioica* mitochondrial genome. CDS are in yellow, rRNAs are in dark blue, and the tRNA-Met gene is in pink. (A) North Sea RNA read coverage (SRR20015061). The average coverage of the genes *rnl* and *rns* is 485,219.8 ± 262,496.2 and 298,863.9 ± 153,147.5 reads per position, respectively. (B) Japan RNA read coverage (SRR1693762, SRR1693765, SRR1693766, SRR1693767). The average coverage of the genes *rnl* and *rns* is 4,440.2 ± 2,033.4 and 66,706.8 ± 44,594.4 reads per position, respectively.

**A.** North Sea RNA coverage

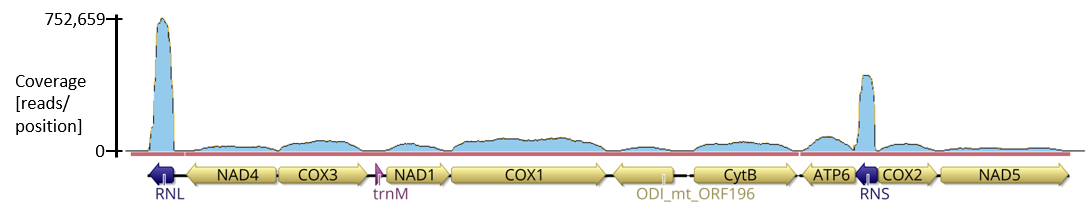

**B.** Japan RNA coverage

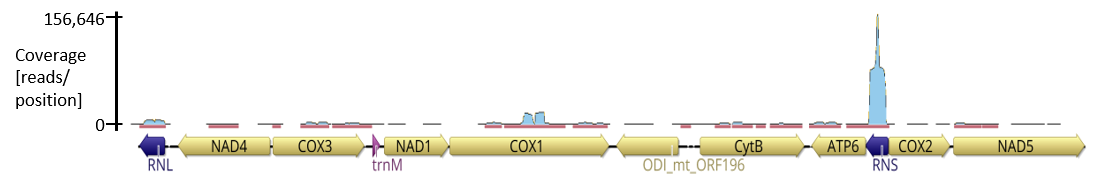

Figure S6: tRNA-Met (UAU)

Secondary structure of tRNA-Met (UAU) predicted using Mitos2.

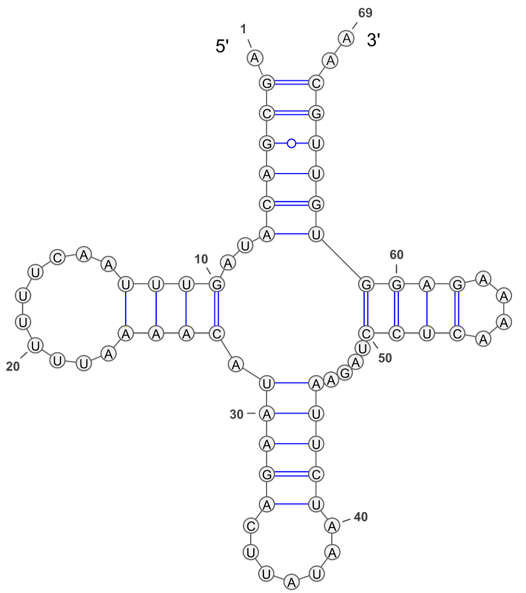

Figure S7: Putative NAD3 protein

Alignment of the putative NAD3 protein of *O. dioica*, encoded in the nuclear genome, with mitochondrially encoded NAD3 protein of animals. The more conserved the alignment position, the darker the amino-acid background. The predicted transmembrane domains and mitochondrial target peptides are indicated in blue and pink, respectively.

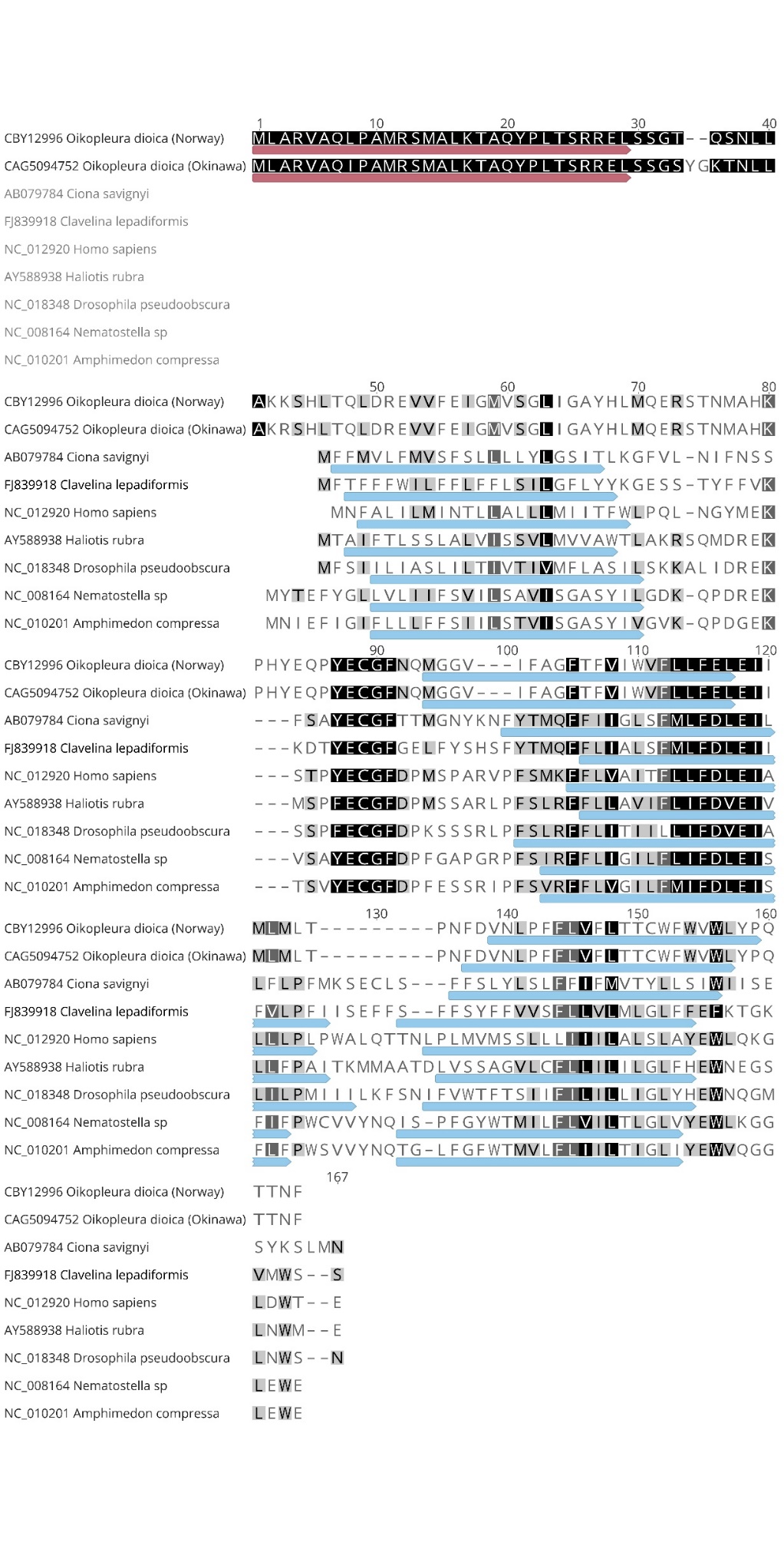

Figure S8A-K: Phylogenetic trees

Phylogenetic tree reconstructed based on mitochondrial protein sequences. Different maximum likelihood models and a Bayesian model were used to reconstruct the phylogenetic trees. Trees were reconstructed twice, either with or without the genus *Salpa* of the Thaliacea class. Branch supports are only indicated for those cases in which the bootstrap support values were lower than 100%.

**A.** mtZOA+F+I+G4 model

-m mtZOA+F+I+G4

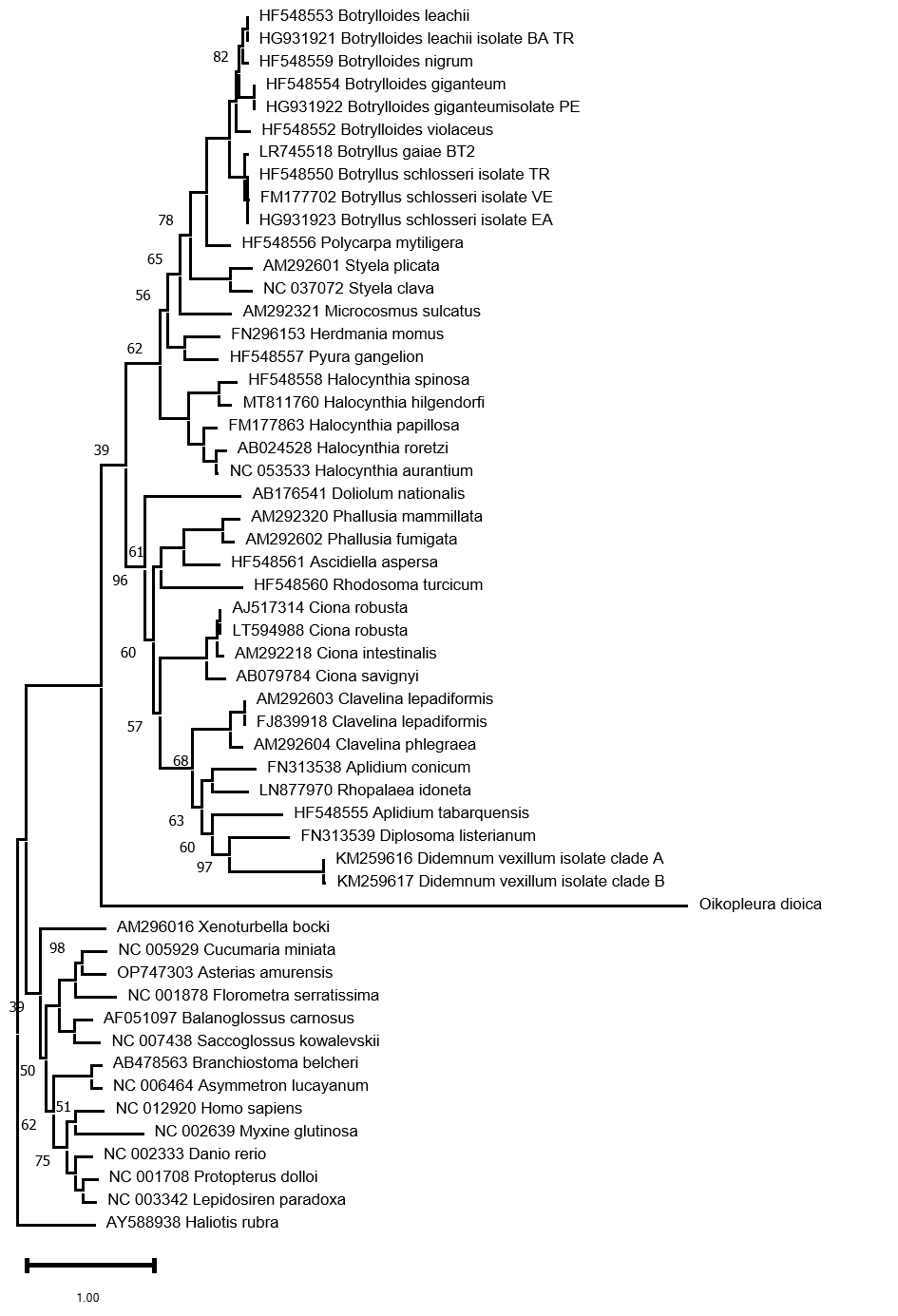

**B.** C10+F model

-mwopt -m C10+F

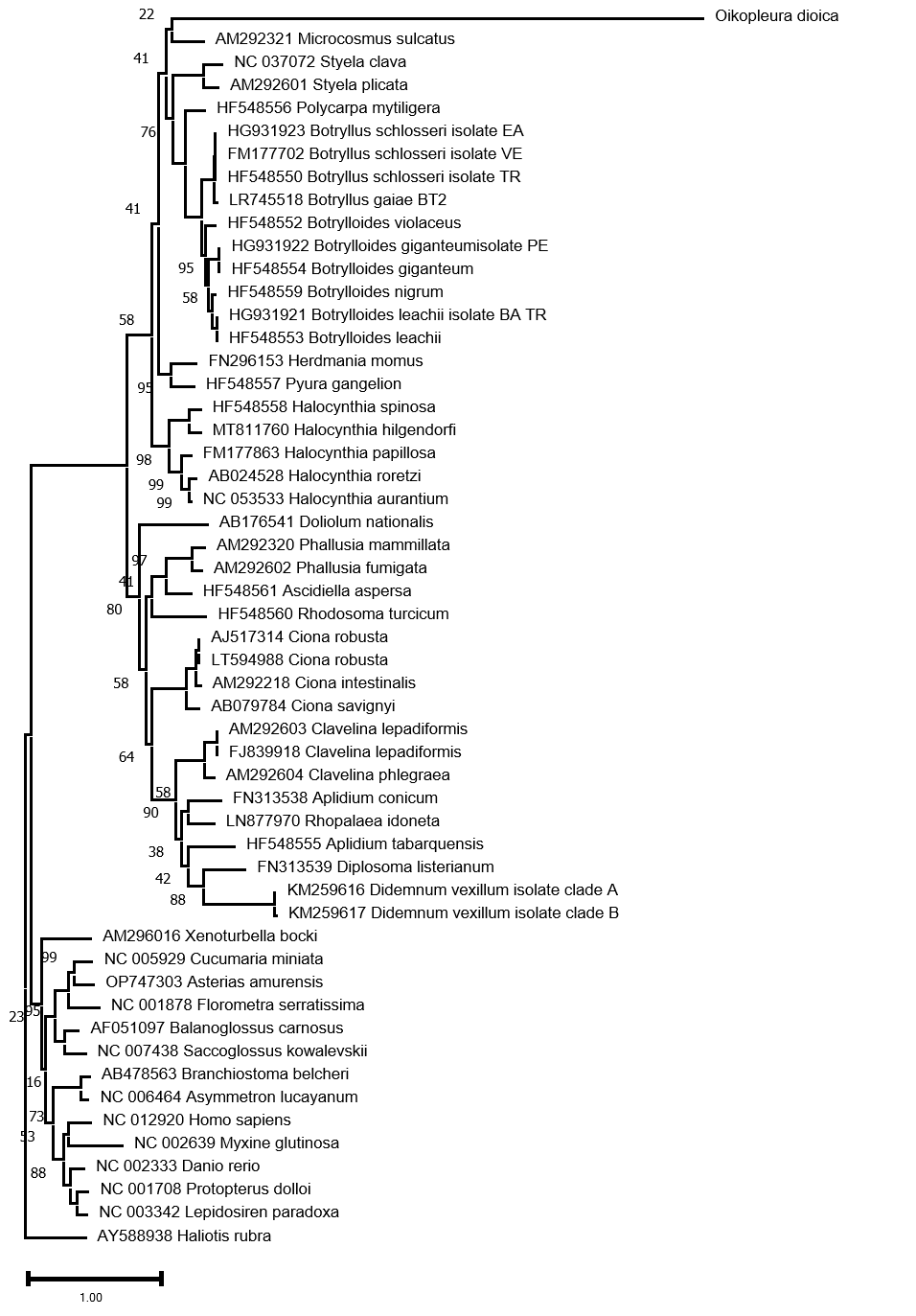

**C.** C20+F model

-mwopt -m C20+F

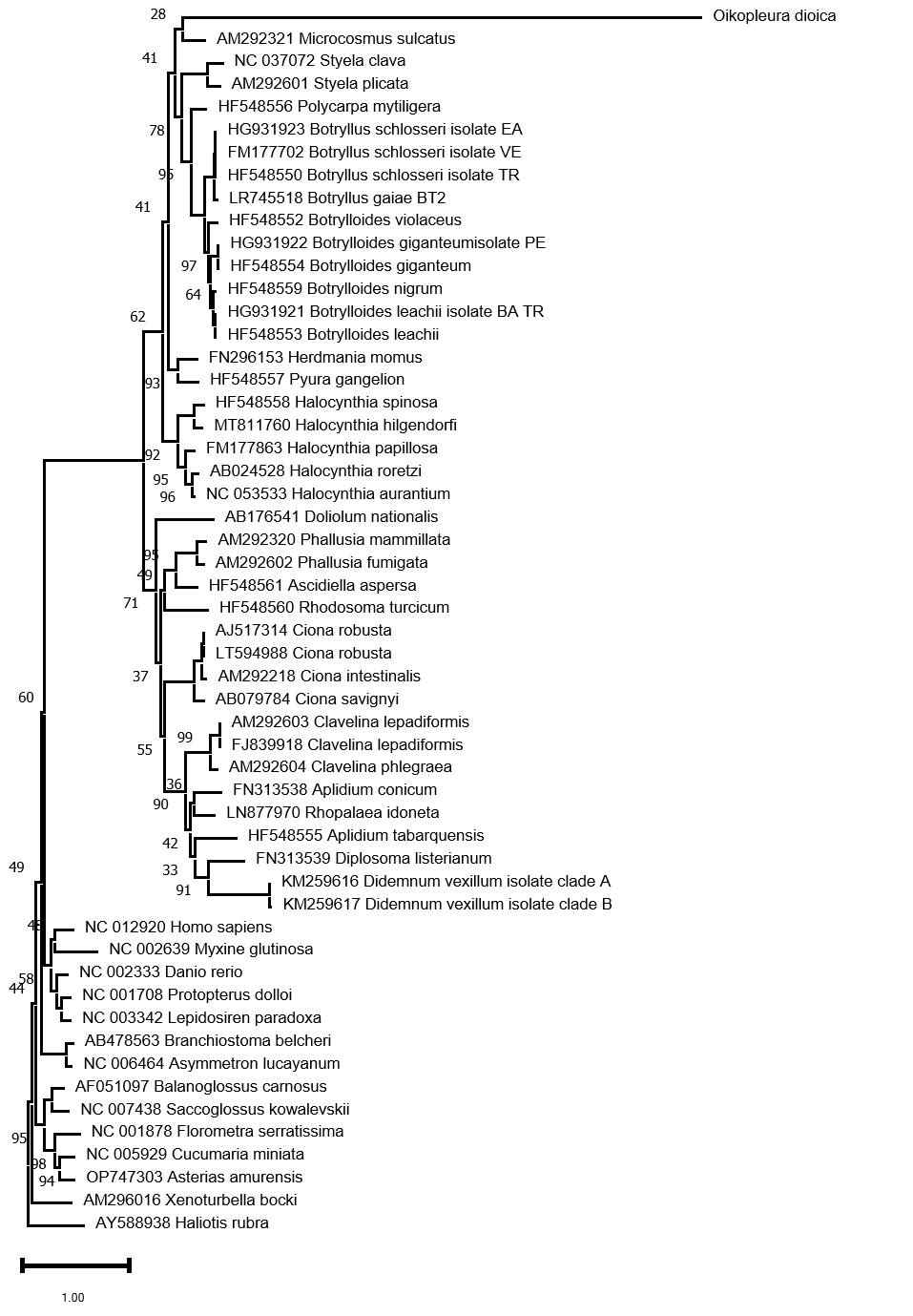

**D.** mtZOA+F+I+G4 model including the *Salpa* genus

-m mtZOA+F+I+G4

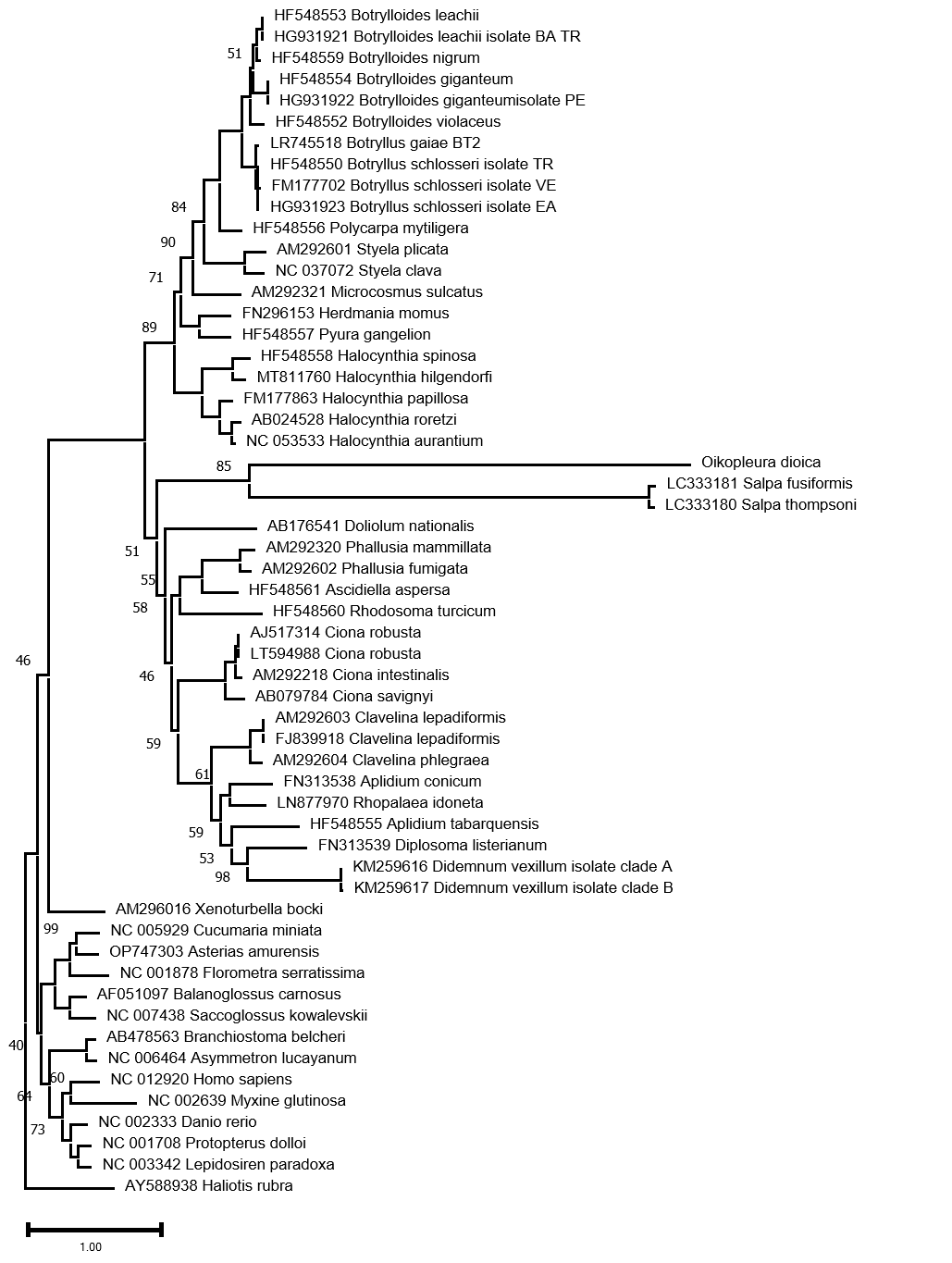

**E.** EHO+F+I+G4 model including the *Salpa* genus

-mwopt -m mtZOA+EHO+F+I+G4

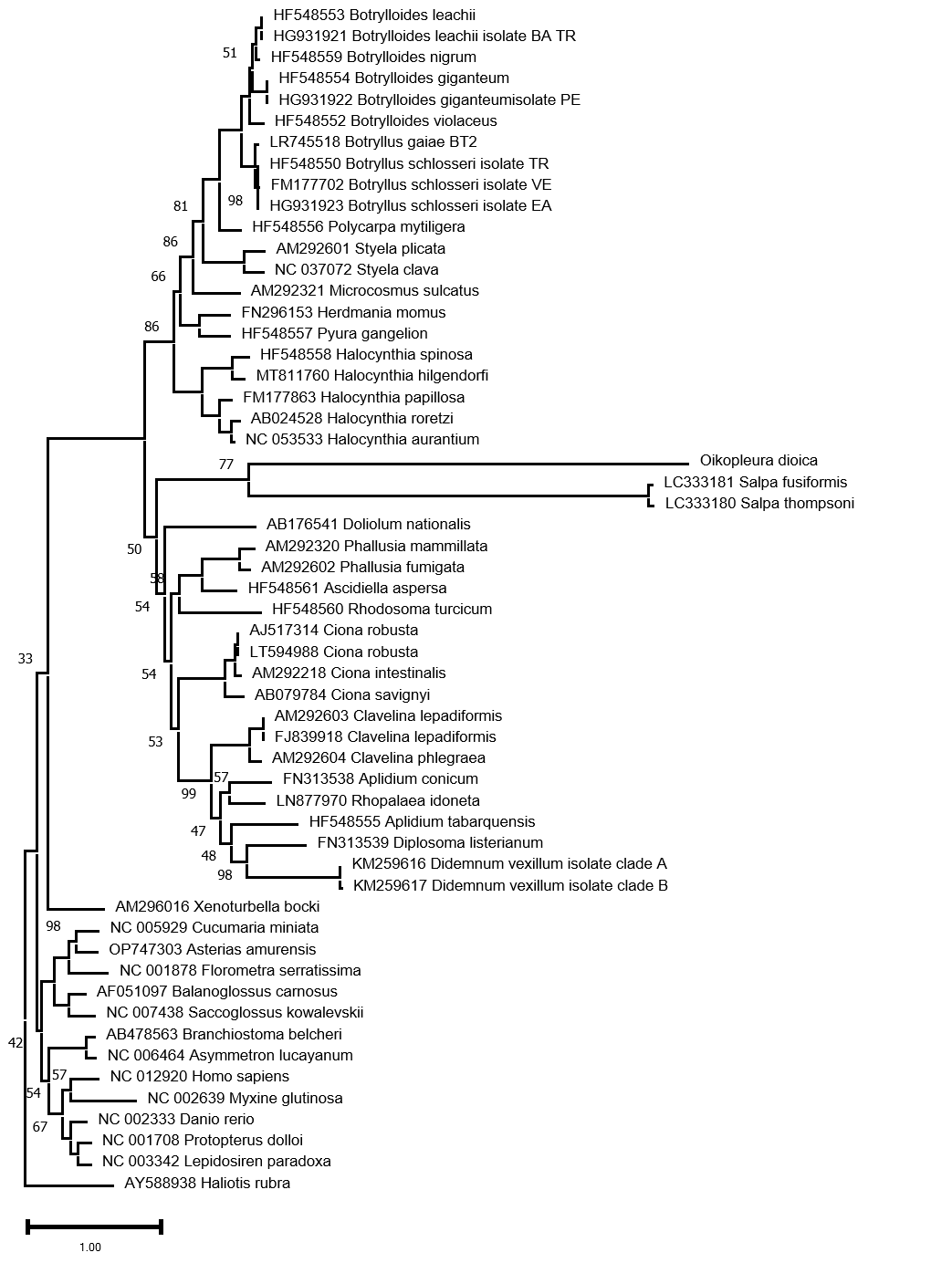

**F.** C10+F model including the *Salpa* genus

-mwopt -m C10+F

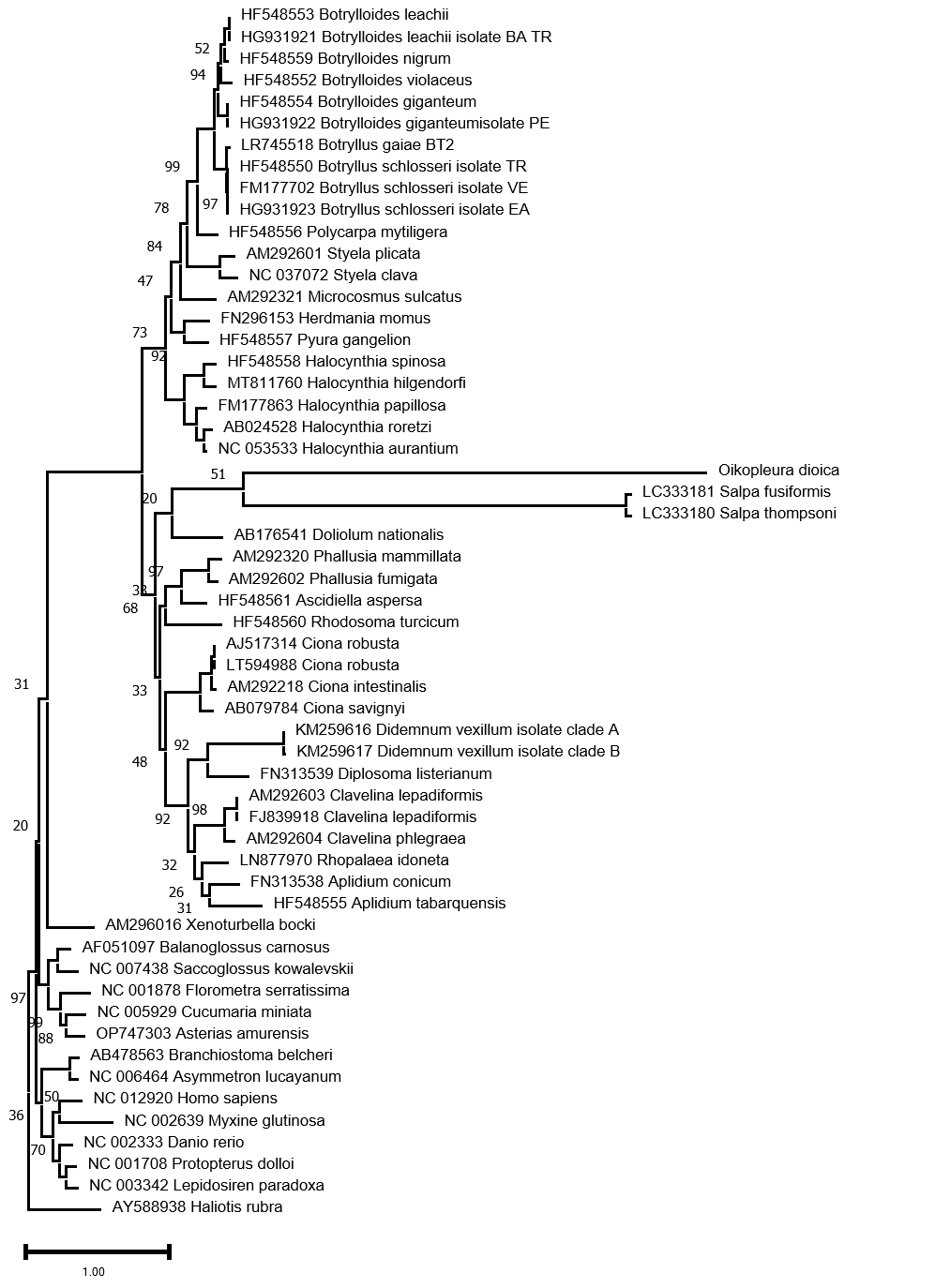

**G.** C20+F model including the *Salpa* genus

-mwopt -m C20+F

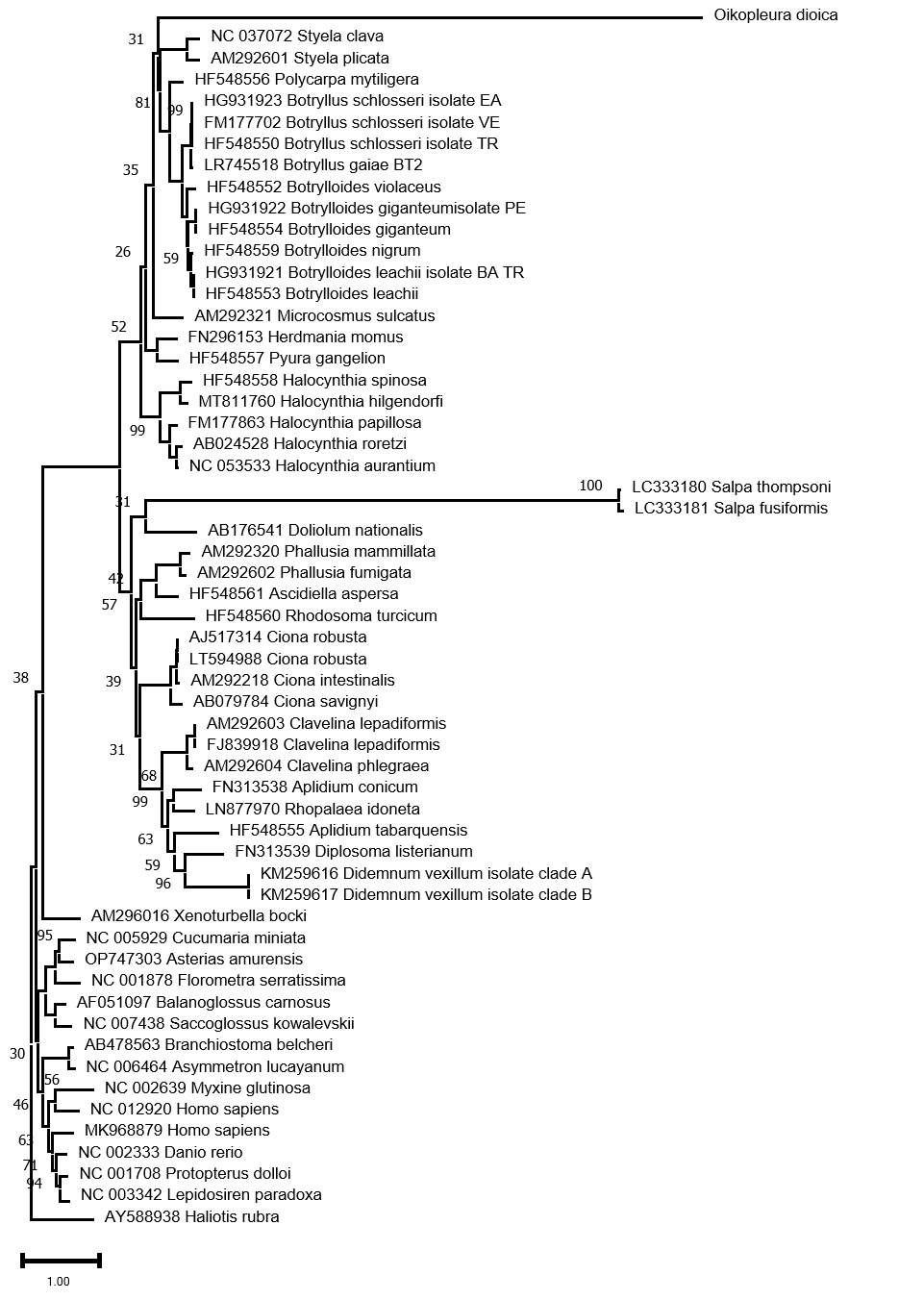

**H.** Phylobase CAT+GTR+G4 model

-cat -gtr -dgam 4
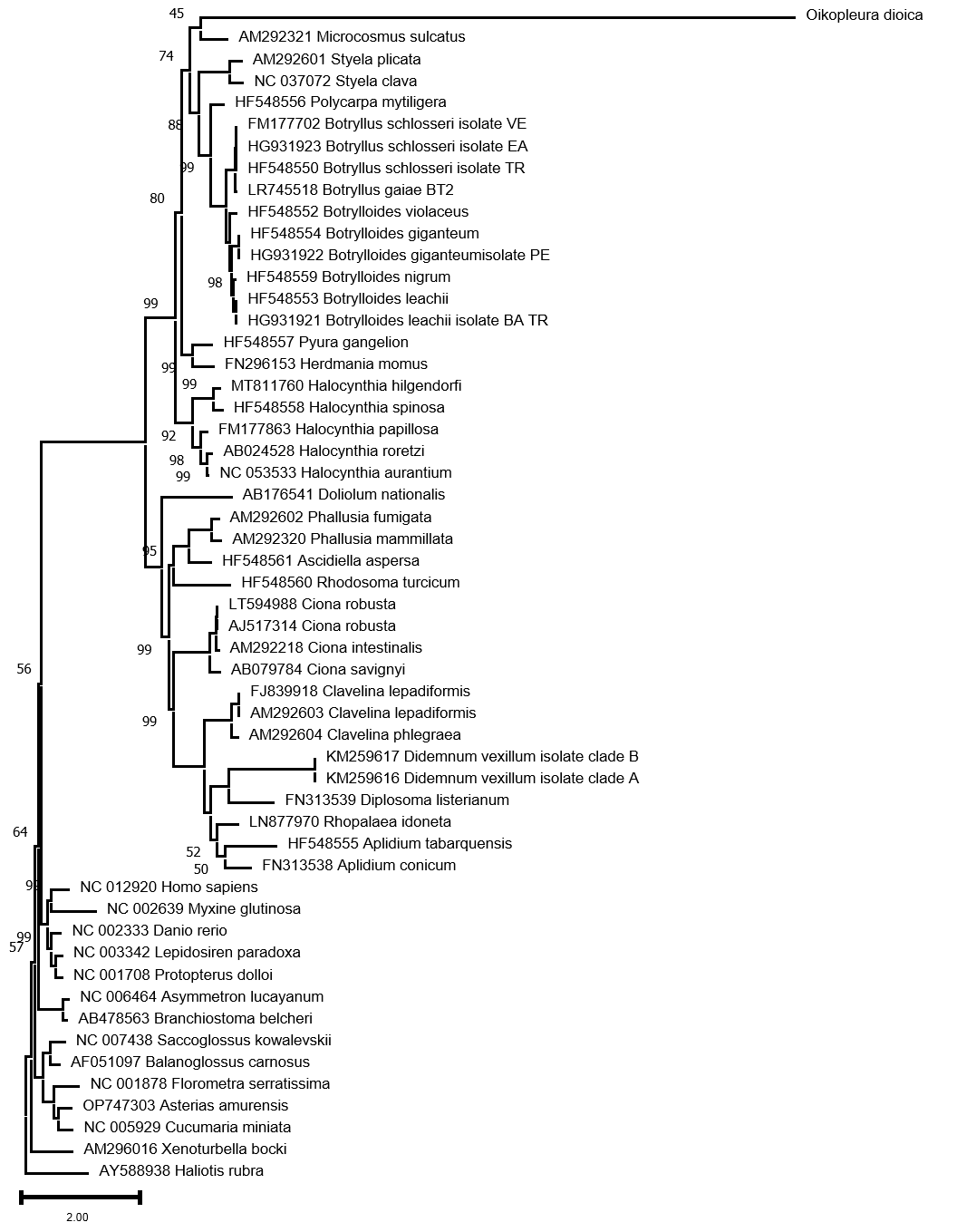

**I.** Phylobase CAT+GTR+G4 model including the *Salpa* genus (chains 124)

-cat -gtr -dgam 4

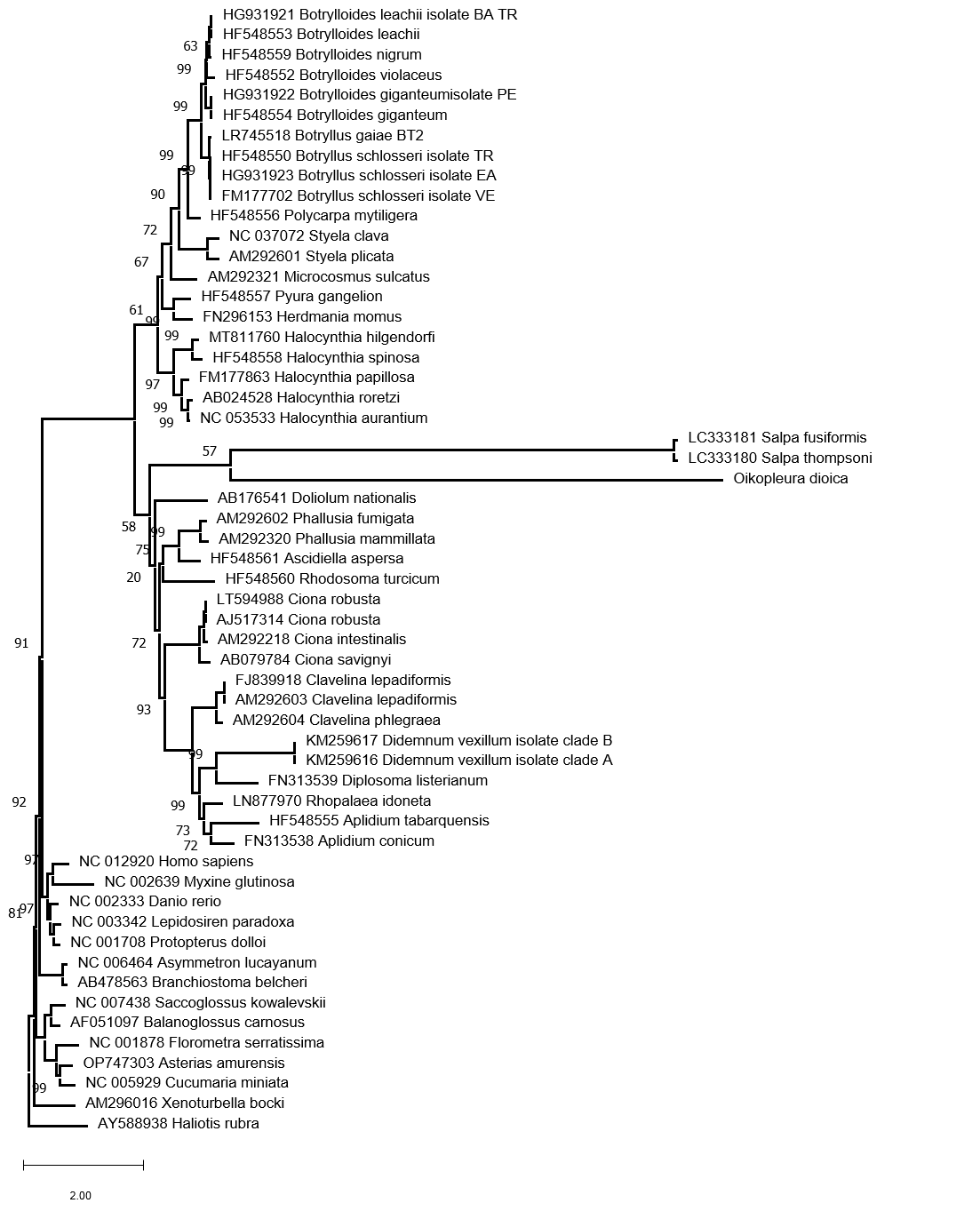

**J.** Phylobase CAT+GTR+G4 model including the *Salpa* genus (chains 123)

-cat -gtr -dgam 4

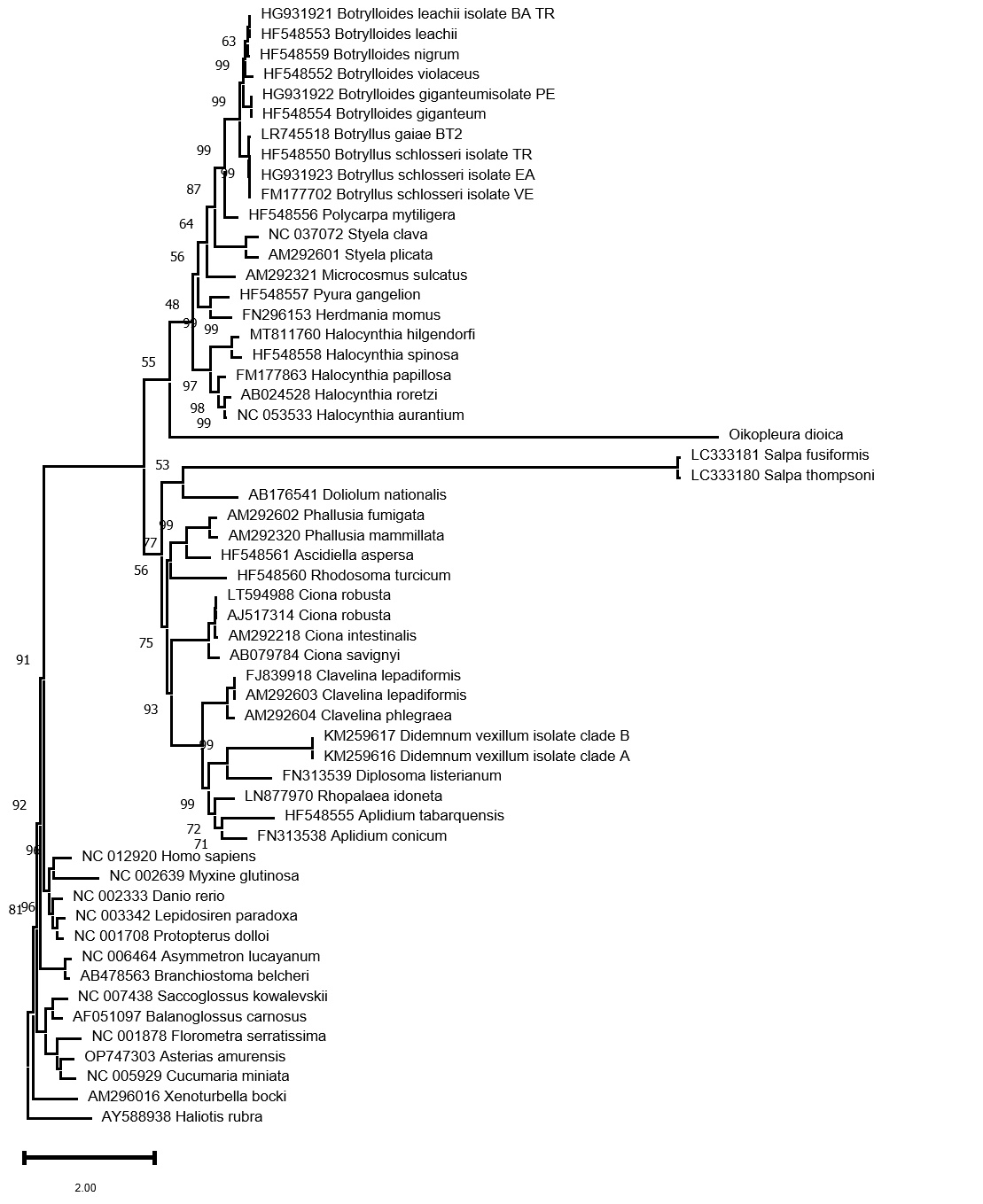

**K.** *O. dioica* phylogenetic positions

Summary of *O. dioica* (in purple) possible positions among tunicates according to the previous phylogenetic trees.

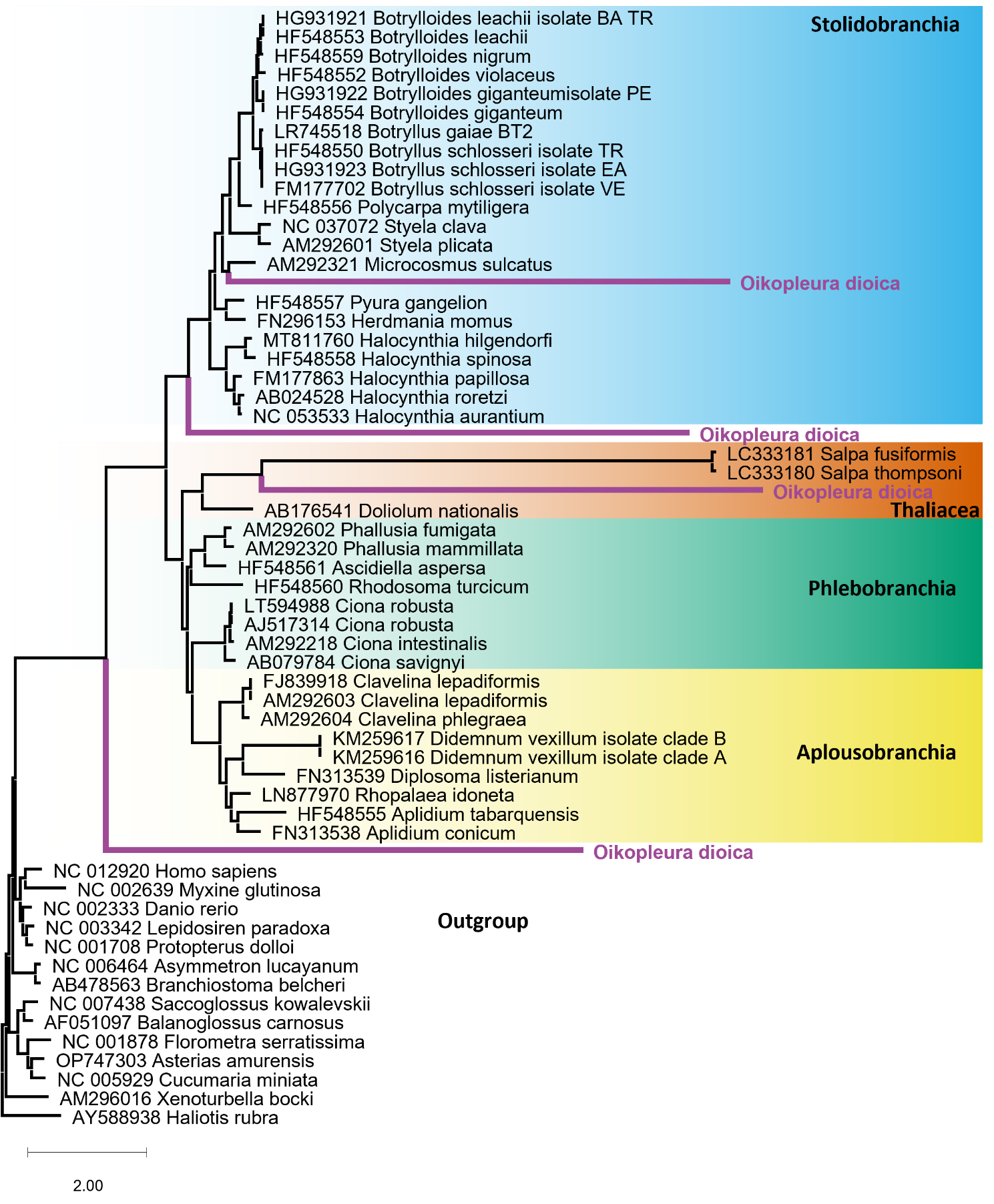

Figure S9: *Cytb* irregular editing site

An irregular editing site in the gene *cytb* that starts with “TTTTTCTT”, which is retained in the transcripts, and is followed by “AACTT”, which is excised (bounded by a purple square). The excised “AACTT” region is present in all DNA reads but is absent in the vast majority of the EST and RNA reads (see Supplementary Table S3 B for accession numbers).

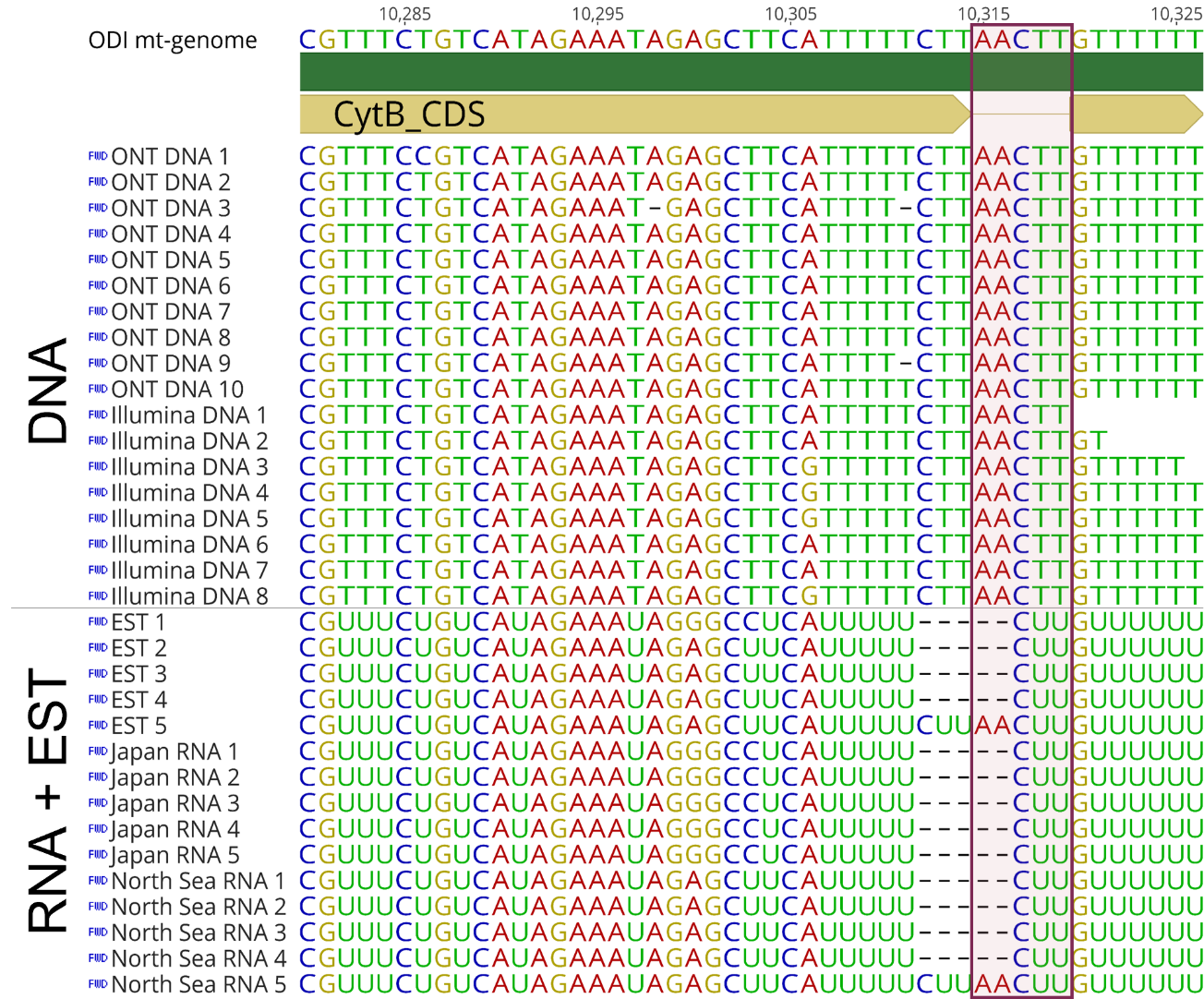

Figure S10: RNA contigs

The genes are in yellow, rRNA are in dark blue, and the tRNA gene is in pink.

(A) *Oikopleura dioica* mitochondrial genes assembled to four contigs based on RNA Illumina reads from Japan (SRR1693762, SRR1693765, SRR1693766, SRR1693767). (B) *Oikopleura dioica* mitochondrial genes assembled to five contigs based on RNA Illumina reads from the North Sea (SRR20015061).

**A.** Japan Contigs:

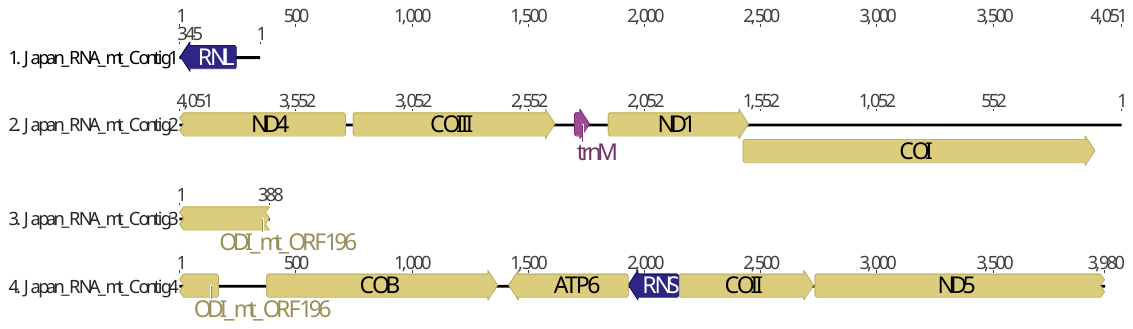

**B.** North Sea Contigs:

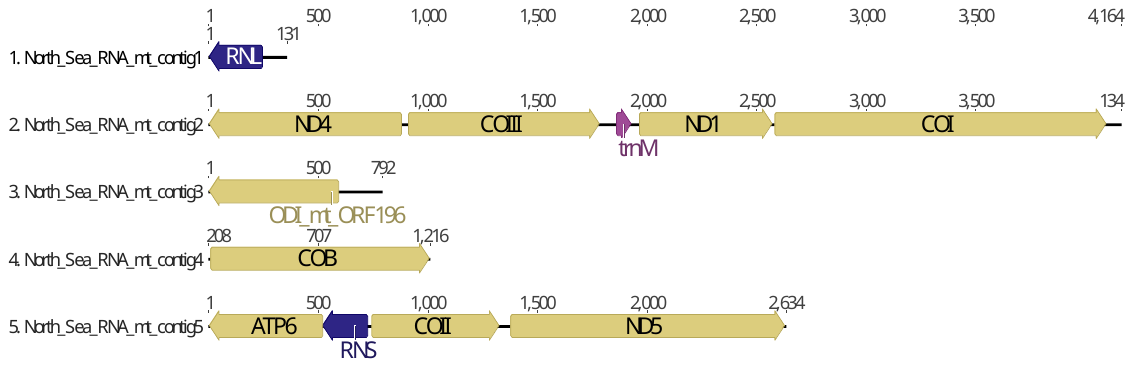

Table S1: PCR primers

Primer pairs were designed to amplify the *O. dioica* mitochondrial genome. Fragment number, primer direction (forward or reverse), primer name, Tm [⁰C], sequence, and approximate length of the mitochondrial PCR products obtained based on gel electrophoresis are provided. The location of the PCR fragments along the final assembled genome is indicated in Figure S1.

| **PCR Frag-ment** | **Primer pairs** | | | | | |  |
| --- | --- | --- | --- | --- | --- | --- | --- |
|  | **Forward** | | | **Reverse** | | |  |
|  |  | **Tm [°C]** | **5'-3'** |  | **Tm [°C]** | **5'-3'** | **Obtained Length [bp]** |
| 1 | 61f | 61.9 | GAAGTCCACGGTCCCTAATATGCAAAAGCA | 31r | 61.9 | CCACAAGCACATAAGAGCAACCCATACACT | 1,400 |
| 2 | 81f | 66.2 | TGTGCTTGTGGACATGGATGGCGGGAG | 9r | 62.4 | GTTCGAGGCCTTGTACGATTTTGAGTGAGC | 2,000 |
| 3 | 9f | 61.3 | GCTCACTCAAAATCGTACAAGGCCTCG | 10r | 62.8 | ACATGTCTGGAGTGGTAACAAATGGGGGAA | 1,500 |
| 4 | 5f | 63.1 | TGGTTGATTCCCCCATTTGTTACCACTCCA | 14r | 62.7 | TAACGACGTGGCATTCCCTCAATTCTCAGT | 1,500 |
| 5 | 14f | 61.5 | CCAAAGGGACCCAAATTTACAATGCTGCT | 14r | 62.7 | TAACGACGTGGCATTCCCTCAATTCTCAGT | 1,200 |
| 6 | 91f | 61 | GGTCTGTACACTATTGGCCTGTATTCTCTGG | 92r | 58.6 | GCCCGAAATGTATAGTCAAAATTACCCTAGTC | 3,000 |
| 7 | 83f | 62.9 | TTTAAAGTACTCACTGCTTCGAGCGCCCTT | 63r | 61.6 | TTTTGCCATGGACTTGTCGAGATTCCTGAT | 3,000 |
| 8 | 72f | 62 | GCGCCCATAACGAAGGTAAGCTCTAACA | 13r | 62.9 | AATGGCGTCACTAGAGGGTTAGCTGAAAGG | 1,400 |
| 9 | 63f | 61.6 | ATCAGGAATCTCGACAAGTCCATGGCAAAA | 41r | 61.8 | AGTGAGAATGCATGTCAGCGGAAGATTGTT | 1,500 |
| 10 | 41f | 60.2 | AACAATCTTCTGCTGACATGCATTCTCACT | 11r | 61.3 | TGTAATAAGCTGGAAACTGGTGTAGGCCTTTC | 3,000 |
| 11 | 11f | 62.4 | CCATAGATAGGCCTACACCAGTTTCCAGCT | 85r | 62.5 | TGGGCCGATATTTAGCACTGATAACCACCT | 3,000 |
| 12 | 7f | 64 | AGAACCTCACGCCTITTACCTGATCCCAAA | 61r | 62 | CTTGCTTTTGCATATTAGGGACCGTGGACT | 2,000 |

Table S2: Characteristics of the poly-A/T stretches

The position, length and direction of the 65 poly-A/T stretches found in the mitochondrial genome.

| **Poly A/T site** | **Position** | | **Length** | **Direction** | **EST/ RNA validation** | **Purine present in the polypyrimidine stretch** |
| --- | --- | --- | --- | --- | --- | --- |
|  | **Start** | **End** |  |  |  |  |
| 1 | 27 | 51 | 25 | reverse | yes | no |
| 2 | 70 | 93 | 24 | reverse | yes | no |
| 3 | 427 | 935 | 509 | forward | no | A |
| 4 | 994 | 1,209 | 216 | reverse | yes | no |
| 5 | 1,293 | 1,456 | 164 | reverse | yes | no |
| 6 | 1,482 | 1,732 | 251 | reverse | yes | no |
| 7 | 1,825 | 2,031 | 207 | reverse | yes | no |
| 8 | 2,289 | 2,410 | 122 | reverse | yes | no |
| 9 | 2,517 | 2,594 | 78 | reverse | yes | no |
| 10 | 2,869 | 2,894 | 26 | reverse | yes | no |
| 11 | 3,094 | 3,307 | 214 | forward | yes | no |
| 12 | 3,484 | 3,712 | 229 | forward | yes | no |
| 13 | 3,943 | 4,170 | 228 | forward | yes | no |
| 14 | 4,296 | 4,423 | 128 | forward | yes | no |
| 15 | 4,573 | 4,588 | 16 | reverse | yes | no |
| 16 | 4,693 | 4,716 | 24 | forward | yes | no |
| 17 | 4,788 | 4,792 | 5 | forward | yes | no |
| 18 | 4,832 | 4,833 | 2 | forward | yes | no |
| 19 | 5,113 | 5,248 | 136 | forward | yes | no |
| 20 | 5,292 | 5,414 | 123 | forward | yes | no |
| 21 | 5,509 | 5,628 | 120 | forward | yes | no |
| 22 | 5,802 | 5,823 | 22 | reverse | yes | no |
| 23 | 5,905 | 5,989 | 85 | forward | yes | no |
| 24 | 6,119 | 6,185 | 67 | forward | yes | no |
| 25 | 6,670 | 6,737 | 68 | forward | yes | no |
| 26 | 6,880 | 6,976 | 97 | forward | yes | no |
| 27 | 7,173 | 7,342 | 170 | forward | yes | no |
| 28 | 7,860 | 7,959 | 100 | reverse | yes | no |
| 29 | 8,008 | 8,146 | 139 | forward | no | no |
| 30 | 8,213 | 8,463 | 251 | forward | no | A |
| 31 | 8,493 | 8,631 | 139 | reverse | yes | no |
| 32 | 8,668 | 8,685 | 18 | reverse | yes | no |
| 33 | 8,753 | 8,932 | 180 | reverse | yes | A |
| 34 | 8,983 | 9,143 | 161 | reverse | yes | no |
| 35 | 9,306 | 9,456 | 151 | reverse | yes | no |
| 36 | 9,526 | 9,711 | 186 | reverse | yes | no |
| 37 | 10,079 | 10,099 | 21 | reverse | no | no |
| 38 | 10,327 | 10,514 | 188 | forward | yes | no |
| 39 | 10,773 | 10,847 | 75 | forward | yes | no |
| 40 | 10,862 | 11,033 | 172 | forward | yes | no |
| 41 | 11,301 | 11,415 | 115 | forward | yes | no |
| 42 | 11,567 | 11,989 | 423 | forward | yes | A |
| 43 | 12,084 | 12,144 | 61 | reverse | no | no |
| 44 | 12,212 | 12,313 | 102 | forward | no | no |
| 45 | 12,346 | 12,490 | 145 | reverse | yes | no |
| 46 | 12,892 | 12,973 | 82 | reverse | yes | no |
| 47 | 13,034 | 13,042 | 9 | reverse | yes | no |
| 48 | 13,226 | 13,242 | 17 | reverse | yes | G |
| 49 | 13,493 | 13,513 | 21 | forward | yes | no |
| 50 | 13,635 | 13,706 | 72 | forward | yes | no |
| 51 | 13,719 | 14,040 | 322 | forward | yes | A |
| 52 | 14,305 | 14,423 | 119 | reverse | yes | no |
| 53 | 14,591 | 14,827 | 237 | forward | yes | A |
| 54 | 14,926 | 15,216 | 291 | forward | yes | A |
| 55 | 15,299 | 15,491 | 193 | forward | yes | no |
| 56 | 15,527 | 15,755 | 229 | forward | yes | no |
| 57 | 15,799 | 16,156 | 358 | forward | yes | no |
| 58 | 16,259 | 16,446 | 188 | forward | yes | no |
| 59 | 16,510 | 16,681 | 172 | forward | yes | no |
| 60 | 16,758 | 16,821 | 64 | forward | yes | no |
| 61 | 16,878 | 17,180 | 303 | forward | yes | no |
| 62 | 17,283 | 17,474 | 192 | forward | yes | no |
| 63 | 17,507 | 17,611 | 105 | forward | yes | no |
| 64 | 17,674 | 17,867 | 194 | forward | yes | no |
| 65 | 18,068 | 18,209 | 142 | forward | yes | no |

Table S3: Read accession numbers

**A.** Accession numbers of the reads used in Figure 2.

| **Read** | **Accession number** |
| --- | --- |
| ONT DNA 1 | 4ca4dc77-52a2-4280-9fd6-856d677e47a0 |
| ONT DNA 2 | f5d7f62b-d2b1-4227-8350-a3ca1dd56369 |
| ONT DNA 3 | e5e101aa-7008-497c-b249-ca9293a2f989 |
| ONT DNA 4 | d4701339-2ec8-45f8-977b-e24b157f4e54 |
| ONT DNA 5 | 3eaf97f3-884d-4da2-b37c-dadd3070275d |
| ONT DNA 6 | 8cc14cff-8cf2-4498-8f13-edf3ba52aa05 |
| ONT DNA 7 | ea687f5d-b13f-4364-8800-7ed179ec008e |
| ONT DNA 8 | 7e3d41ef-75d7-461a-806e-cb00c85f94f6 |
| Illumina DNA 1 | HISEQ295CDW7RANXX211051903022010 |
| Illumina DNA 2 | HISEQ295CDW7RANXX21107717967260 |
| Illumina DNA 3 | HISEQ295CDW7RANXX211042111559438 |
| Illumina DNA 4 | HISEQ295CDW7RANXX211021071861725 |
| Illumina DNA 5 | HISEQ295CDW7RANXX21102986245095 |
| Illumina DNA 6 | HISEQ295CDW7RANXX211041214171042 |
| Illumina DNA 7 | HISEQ295CDW7RANXX211091081995773 |
| Illumina DNA 8 | HISEQ295CDW7RANXX211021094876280 |
| EST 1 | KT0AAA3YI20.CONTIG |
| EST 2 | KT0AAA3YK22.CONTIG |
| EST 3 | KT0AAA6YG11.CONTIG |
| EST 4 | KT0AAA8YB09.CONTIG |
| EST 5 | KT0AAA16YL02.CONTIG |
| Japan RNA 1 | SRR1693767.3203366.2 FCC3HP5ACXX412051250266529# |
| Japan RNA 2 | SRR1693767.3203366.2 FCC3HP5ACXX412051250266529# |
| Japan RNA 3 | SRR1693765.7461688.1 FCC3HP5ACXX421052098352519# |
| Japan RNA 4 | SRR1693762.739614.2 FCC3HP5ACXX311031409387534# |
| Japan RNA 5 | SRR1693762.8946134.2 FCC3HP5ACXX32110784833559# |
| North Sea RNA 1 | SRR20015061.16153347 16153347/1 |
| North Sea RNA 2 | SRR20015061.18112952 18112952/2 |
| North Sea RNA 3 | SRR20015061.18804463 18804463/2 |
| North Sea RNA 4 | SRR20015061.19119916 19119916/1 |
| North Sea RNA 5 | SRR20015061.21468084 21468084/2 |

**B.** Accession numbers of the reads used in Figure S9.

| **Read** | **Accession number** |
| --- | --- |
| ONT DNA 1 | defcdc6b-d33b-4529-aa4a-7c773228075e |
| ONT DNA 2 | 73bca372-6389-4006-be53-b0962ef1431c |
| ONT DNA 3 | c5005b75-97a5-4ff3-bec9-84660a8a742d |
| ONT DNA 4 | 67128097-8267-476f-9dab-95d1b4165661 |
| ONT DNA 5 | 91b52ea3-4d1f-48cf-bddb-8f5ecf44a4d0 |
| ONT DNA 6 | 694824fa-f084-4230-9081-802205cd5527 |
| ONT DNA 7 | 6fd36deb-b7b6-4e77-87a4-c12e908b8706 |
| ONT DNA 8 | 08151529-7e49-4be2-adcd-12875bdd2061 |
| ONT DNA 9 | 7f014cac-033c-4a7a-8694-f0f5a4a6fcce |
| ONT DNA 10 | 844f513d-6db2-4cd3-8d09-adf4cbba1a4f |
| ONT DNA 11 | a14ed876-84fe-4114-a361-eec831888995 |
| ONT DNA 12 | 1f8862ac-c482-4131-afad-6100cf458bf4 |
| Illumina DNA 1 | HISEQ295CDW7RANXX211011963614865 |
| Illumina DNA 2 | HISEQ295CDW7RANXX2110248882227 |
| Illumina DNA 3 | HISEQ295CDW7RANXX21101193887642 |
| Illumina DNA 4 | HISEQ295CDW7RANXX21102665080900 |
| Illumina DNA 5 | HISEQ295CDW7RANXX211031462650996 |
| Illumina DNA 6 | HISEQ295CDW7RANXX211021465356572 |
| Illumina DNA 7 | HISEQ295CDW7RANXX21102118342298 |
| Illumina DNA 8 | HISEQ295CDW7RANXX211011360075946 |
| Illumina DNA 9 | HISEQ295CDW7RANXX211061724944422 |
| Illumina DNA 10 | HISEQ295CDW7RANXX21110114325671 |
| EST 1 | KT0AAA12YC07.CONTIG |
| EST 2 | KT0AAA134YM10RM1.SCF |
| EST 3 | KT0AAA148YO11RM1.SCF |
| EST 4 | KT0AAA24YI17.CONTIG |
| EST 5 | KT0AAA5YF24.CONTIG |
| Japan RNA 1 | SRR1693765.2471225.2 FCC3HP5ACXX41205450813128# |
| Japan RNA 2 | SRR1693765.5471485.2 FCC3HP5ACXX413071490223242# |
| Japan RNA 3 | SRR1693762.1039485.2 FCC3HP5ACXX311081043271679# |
| Japan RNA 4 | SRR1693762.5533894.2 FCC3HP5ACXX31307679736771# |
| Japan RNA 5 | SRR1693762.8615839.2 FCC3HP5ACXX3211372486793# |
| North Sea RNA 1 | SRR20015061.14112058 14112058/1 |
| North Sea RNA 2 | SRR20015061.17631641 17631641/2 |
| North Sea RNA 3 | SRR20015061.1439979 1439979/1 |
| North Sea RNA 4 | SRR20015061.2325314 2325314/2 |
| North Sea RNA 5 | SRR20015061.24606496 24606496/2 |
| North Sea RNA 6 | SRR20015061.25754014 25754014/1 |

Table S4: Sequences included in the phylogenetic analysis

Accession numbers of the 55 complete mitochondrial genomes used to construct the phylogenetic tree. Of note the 41 tunicate sequences were used to compute the average gene lengths provided in Table 1.

| **Classification** | **Species** | **Association Number** |
| --- | --- | --- |
| Thaliacea | *Doliolum nationalis* | AB176541 |
|  | *Salpa fusiformis* | LC333181 |
|  | *Salpa thompsoni* | LC333180 |
| Stolidobranchia | *Botrylloides giganteum* | HF548554; isolate PE HG931922 |
|  | *Botrylloides leachii* | HF548553; isolate BA TR HG931921 |
|  | *Botrylloides nigrum* | HF548559 |
|  | *Botrylloides violaceus* | HF548552 |
|  | *Botryllus gaiae* | isolate BT2 LR745518 |
|  | *Botryllus schlosseri* | isolate VE FM177702; isolate TR HF548550; isolate EA HG931923 |
|  | *Halocynthia aurantium* | NC 53533 |
|  | *Halocynthia hilgendorfi* | MT811760 |
|  | *Halocynthia papillosa* | FM177863 |
|  | *Halocynthia roretzi* | AB024528 |
|  | *Halocynthia spinosa* | HF548558 |
|  | *Herdmania momus* | FN296153 |
|  | *Microcosmus sulcatus* | AM292321 |
|  | *Polycarpa mytiligera* | HF548556 |
|  | *Pyura gangelion* | HF548557 |
|  | *Styela clava* | NC 37072 |
|  | *Styela plicata* | AM292601 |
| Phlebobranchia | *Ascidiella aspersa* | HF548561 |
|  | *Ciona intestinalis* | AM292218 |
|  | *Ciona robusta* | AJ517314, LT594988 |
|  | *Ciona savignyi* | AB079784 |
|  | *Phallusia fumigata* | AM292602 |
|  | *Phallusia mammillata* | AM292320 |
|  | *Rhodosoma turcicum* | HF548560 |
| Aplousobranchia | *Aplidium conicum* | FN313538 |
|  | *Aplidium tabarquensis* | HF548555 |
|  | *Clavelina lepadiformis* | AM292603, FJ839918 |
|  | *Clavelina phlegraea* | AM292604 |
|  | *Didemnum vexillum* | isolate clade A KM259616; isolate clade B KM259617 |
|  | *Diplosoma listerianum* | FN313539 |
|  | *Rhopalaea idoneta* | LN877970 |
| Cephalochordata | *Asymmetron lucayanum* | NC_006464 |
|  | *Branchiostoma belcheri* | AB478563 |
| Vertebrata | *Danio rerio* | NC_002333 |
|  | *Homo sapiens* | NC_012920 |
|  | *Lepidosiren paradoxa* | NC_003342 |
|  | *Myxine glutinosa* | NC 002639 |
|  | *Protopterus dolloi* | NC_001708 |
| Echinodermata | *Asterias amurensis* | OP747303 |
|  | *Cucumaria miniata* | NC_005929 |
|  | *Florometra serratissima* | NC_001878 |
| Hemichordata | *Balanoglossus carnosus* | AF051097 |
|  | *Saccoglossus kowalevskii* | NC_007438 |
| Spiralia | *Haliotis rubra* | AY588938 |
| Xenacoelomorpha | *Xenoturbella bocki* | AM296016 |

Table S5: Aminoacyl-tRNA synthetase (aaRS) accession numbers

Accession numbers of the aaRS sequences analyzed in Figure 3. Single eukaryotic (Euk): single copy aaRSs that function both in the cytosol and mitochondria. Other aaRSs are either expressed only in the cytosol (cytosolic) or in the mitochondrion (mitochondrial). In pink aaRSs presence, in white absence, in blue the aaRS-Gly, for which one cytosolic and one mitochondrial gene are present *in O. dioica.*

|  | **AA** | *Oikopleura dioica*  Norway  GCA_000209535 | *Oikopleura dioica*  *O*kinawa  GCA_907165135 | *Ciona intestinalis*  GCA_000224145 | *Styela clava*  GCF_013122585 | *Homo sapiens*  GCF_000001405 |
| --- | --- | --- | --- | --- | --- | --- |
| Single Euk. | **A** | CBY23722 | CAG5086037 | XP_002125465 | XP_039260162 | NP_001596 |
|  | **G** |  |  | XP_026689616 | XP_039267458 | NP_002038 |
|  | **H** | CBY19939 | CAG5098982 | XP_002125799 | XP_039265303 | NP_002100 |
|  | **Q** | CBY24607 | CAG5097438 | XP_002131102 | XP_039253901 | NP_005042 |
| Cytosolic | **G** | CBY07672 | CAG5110020 |  |  |  |
|  | **C** | CBY17967 | CAG5114156 | XP_026692167 | XP_039251784 | NP_001014437 |
|  | **D** | CBY13303 | CAG5097808 | XP_002125902 | XP_039267320 | NP_001340 |
|  | **E/P** | CBY12760 | CAG5087613 | XP_002126649 | XP_039268249 | NP_004437 |
|  | **Fa** | CBY20270 | CAG5113901 | XP_002128146 | XP_039274465 | NP_004452 |
|  | **Fb** | CBY22307 | CAG5105684 | XP_002122110 | XP_039249843 | NP_005678 |
|  | **I** | CBY22011 | CAG5105485 | XP_026690541 | XP_039263025 | NP_001365507 |
|  | **K** | CBY15807 | CAG5086875 | XP_002131983 | XP_039249882 | NP_005539 |
|  | **L** | CBY19027 | CAG5091338 | XP_009857630 | XP_039265438 | NP_064502 |
|  | **M** | CBY22485 | CAG5113570 | XP_002128793 | XP_039252310 | NP_004981 |
|  | **N** | CBY09464 | CAG5096130 | XP_002128269 | XP_039266968 | NP_004530 |
|  | **R** | CBY20749 | CAG5113835 | XP_002129614 | XP_039257488 | NP_002878 |
|  | **S** | CBY22081 | CAG5107347 | XP_002127248 | XP_039260596 | NP_006504 |
|  | **T** | CBY19737 | CAG5097418 | XP_009862171 | XP_039248388 | NP_001245366 |
|  | **V** | CBY11536 | CAG5080332 | XP_026693603 | XP_039269212 | XP_054184482 |
|  | **W** | CBY24158 | CAG5105750 | XP_002122401 | XP_039254452 | NP_998810 |
|  | **Y** | CBY24854 | CAG5087034 | XP_002129278 | XP_039272029 | NP_003671 |
| Mitochondrial | **G** | CBY18832 | CAG5105817 |  |  |  |
|  | **C** |  |  | XP_026692488 | XP_039265970 | NP_078813 |
|  | **D** |  |  | XP_026692423 | XP_039255484 | NP_060592 |
|  | **E** |  |  | XP_002127194 | XP_039270239 | NP_001077083 |
|  | **F** | CBY08952 | CAG5096278 | XP_009857978 | XP_039273503 | NP_001305801 |
|  | **I** |  |  | XP_002131649 | XP_039259013 | NP_060530 |
|  | **K** |  |  |  |  |  |
|  | **L** |  |  | XP_002129281 | XP_039249842 | NP_001355192 |
|  | **M** | CBY19815 | CAG5097262 | XP_002126600 | XP_039266187 | NP_612404 |
|  | **N** |  |  | XP_002119984 | XP_039251824 | NP_078954 |
|  | **P** |  |  | XP_009858012 | XP_039248001 | NP_689481 |
|  | **R** |  |  | XP_002126846 | XP_039257488 | NP_064716 |
|  | **S** |  |  | XP_002123180 | XP_039262234 | NP_060297 |
|  | **T** |  |  |  |  |  |
|  | **V** |  |  |  |  | NP_065175 |
|  | **W** |  |  | XP_026690654 | XP_039266839 | NP_056651 |
|  | **Y** |  |  | XP_018668493 | XP_039262919 | NP_001035526 |

Table S6: Codon usage statistics

The initiation codons are in orange, stop codons are in blue, and the most frequent codon, “TTT”, is in pink.

| **Codon** | **AA** | **% of AA** | **Freq** | **% of codon** |  |  | **Codon** | **AA** | **% of AA** | **Freq** | **% of codon** |
| --- | --- | --- | --- | --- | --- | --- | --- | --- | --- | --- | --- |
| GCA | A | 16.67% | 18 | 1.06% |  |  | ATC | M | 0.43% | 1 | 0.06% |
| GCC | A | 33.33% | 36 | 2.12% |  |  | ATT | M | 1.72% | 4 | 0.24% |
| GCG | A | 0.93% | 1 | 0.06% |  |  | AAC | N | 40.82% | 40 | 2.36% |
| GCT | A | 49.07% | 53 | 3.13% |  |  | AAT | N | 59.18% | 58 | 3.42% |
| TGC | C | 41.18% | 21 | 1.24% |  |  | CCA | P | 31.75% | 20 | 1.18% |
| TGT | C | 58.82% | 30 | 1.77% |  |  | CCC | P | 7.94% | 5 | 0.29% |
| GAC | D | 48.28% | 28 | 1.65% |  |  | CCG | P | 3.17% | 2 | 0.12% |
| GAT | D | 51.72% | 30 | 1.77% |  |  | CCT | P | 57.14% | 36 | 2.12% |
| GAA | E | 62.50% | 40 | 2.36% |  |  | CAA | Q | 80.56% | 29 | 1.71% |
| GAG | E | 37.50% | 24 | 1.42% |  |  | CAG | Q | 19.44% | 7 | 0.41% |
| TTC | F | 28.37% | 80 | 4.72% |  |  | CGA | R | 11.11% | 4 | 0.24% |
| TTT | F | 71.63% | 202 | 11.91% |  |  | CGC | R | 8.33% | 3 | 0.18% |
| AGA | G | 59.34% | 143 | 8.43% |  |  | CGG | R | 11.11% | 4 | 0.24% |
| AGG | G | 14.52% | 35 | 2.06% |  |  | CGT | R | 69.44% | 25 | 1.47% |
| GGA | G | 13.69% | 33 | 1.95% |  |  | AGC | S | 15.10% | 37 | 2.18% |
| GGC | G | 1.66% | 4 | 0.24% |  |  | AGT | S | 11.84% | 29 | 1.71% |
| GGG | G | 4.56% | 11 | 0.65% |  |  | TCA | S | 18.78% | 46 | 2.71% |
| GGT | G | 6.22% | 15 | 0.88% |  |  | TCC | S | 13.88% | 34 | 2.00% |
| CAC | H | 35.94% | 23 | 1.36% |  |  | TCG | S | 3.67% | 9 | 0.53% |
| CAT | H | 64.06% | 41 | 2.42% |  |  | TCT | S | 36.73% | 90 | 5.31% |
| ATC | I | 23.39% | 40 | 2.36% |  |  | ACA | T | 40.67% | 61 | 3.60% |
| ATT | I | 76.61% | 131 | 7.72% |  |  | ACC | T | 21.33% | 32 | 1.89% |
| AAA | K | 72.22% | 65 | 3.83% |  |  | ACG | T | 7.33% | 11 | 0.65% |
| AAG | K | 27.78% | 25 | 1.47% |  |  | ACT | T | 30.67% | 46 | 2.71% |
| CTA | L | 20.94% | 71 | 4.19% |  |  | GTA | V | 45.00% | 90 | 5.31% |
| CTC | L | 2.36% | 8 | 0.47% |  |  | GTC | V | 11.00% | 22 | 1.30% |
| CTG | L | 7.37% | 25 | 1.47% |  |  | GTG | V | 19.50% | 39 | 2.30% |
| CTT | L | 11.50% | 39 | 2.30% |  |  | GTT | V | 24.50% | 49 | 2.89% |
| TTA | L | 36.58% | 124 | 7.31% |  |  | TGA | W | 34.38% | 22 | 1.30% |
| TTG | L | 21.24% | 72 | 4.25% |  |  | TGG | W | 65.63% | 42 | 2.48% |
| ATA | M | 55.79% | 130 | 7.67% |  |  | TAC | Y | 44.70% | 59 | 3.48% |
| ATG | M | 40.77% | 95 | 5.60% |  |  | TAT | Y | 55.30% | 73 | 4.30% |
| GTG | M | 0.43% | 1 | 0.06% |  |  | TAA | * | 58.70% | 27 | 1.59% |
| TTG | M | 0.86% | 2 | 0.12% |  |  | TAG | * | 41.30% | 19 | 1.12% |
